# The bifunctional dynamin-like GTPase switch DynAB modulates both vegetative and sporulation cell division in *Streptomyces*

**DOI:** 10.1101/2025.06.23.661029

**Authors:** Meng Liu, Hanlei Zhang, Zhaoxu Jiang, Yanhong Zeng, Zhilong Zhao, Yanping Zhu, Peipei Zhang, Xiuhua Pang

## Abstract

As part of their complex developmental lifecycle, the multicellular filamentous bacteria *Streptomyces* have evolved two modes of cell division distinct from binary fission. In vegetative cell division, irregularly spaced cross-walls form within vegetative hyphae, whereas in sporogenic hyphae, regularly spaced sporulation septa form synchronously, leading to spore formation. However, the mechanisms controlling these processes largely remain unknown. The bacterial dynamin-like protein pair DynAB was previously reported to be a component of sporulation septa. Here, we reveal that DynA and DynB have critical roles in both cell division modes, regulating the formation of sporulation septa and cross-walls, with the *Streptomyces* cell division protein SsgB modulating their activities. *dynAB* deletion and overexpression, respectively, increased and abrogated cross-wall formation in vegetative hyphae, and *Streptomyces* DynAB also inhibited cell division in *Escherichia coli*, resulting in formation of unicellular filamentous cells and suggesting an ancient and conserved function for these proteins. Notably, SsgB could relieve the inhibition of cell division by DynAB in both *Streptomyces* and *E. coli*. In contrast to DynAB, SsgB overexpression generated spore-like compartments in vegetative hyphae, a phenomenon that required disruption of the DynAB complex. Fluorescent tags revealed dynamic localization of DynAB during development, and further analyses indicated that, in sporogenic hyphae, the timing of DynAB expression, their GTP-binding activity, and interaction with SsgB were associated with the synchronous initiation of sporulation septation. Our findings establish DynAB as an integrator of spatiotemporal cues in bacterial multicellularity and provide insights into the evolution of complex cell cycle regulation in prokaryotes.

**Significance:** The evolution of multicellularity in bacteria involved the development of complex mechanisms to spatially and temporally coordinate cell division. *Streptomyces*, renowned for their production of bioactive secondary metabolites, exemplify bacterial complexity. This study reveals how *Streptomyces* employ the dynamin-like proteins DynAB as a bifunctional molecular switch, enabling transition between vegetative and reproductive growth through integrated interactions with developmentally regulated proteins and the availability of the energy supplier GTP. Importantly, DynAB could repress cell division in both *Streptomyces* and *Escherichia coli*, suggesting an evolutionary origin of these proteins prior to bacterial multicellularity. Our work reveals the dual function and switch mechanism of DynAB in *Streptomyces*, providing insights into the emergence of dynamic cell cycle control in prokaryotes.

## Introduction

In unicellular bacteria, cell division is epitomized by binary fission, a meticulously orchestrated event where cells undergo symmetrical splitting at the mid-cell position. Central to this spatial precision is the dynamic assembly of the Z-ring, which is a contractile structure guided by FtsZ polymerization and which acts as the core of the divisome (1). The divisome coordinates membrane constriction and septal peptidoglycan synthesis by resolving checkpoint barriers through multivalent protein interactions (2). Compared to unicellular bacteria, the filamentous actinomycetes and cyanobacteria have evolved complex division types to coordinate their multicellular developmental programs (3–5). Bacteria of the actinomycetes genus *Streptomyces* undergo two distinct modes of cell division: vegetative cross-wall formation, which supports hyphal network expansion, and sporulation-specific septation, which enables synchronized spore production (6). Concurrently with the transition from vegetative growth to reproduction growth, *Streptomyces* synthesize diverse secondary metabolites, including clinically vital antibiotics, through metabolic pathways tightly coupled to developmental transitions (6). Despite the ecological and biomedical significance of these processes, understanding the molecular mechanisms governing the spatiotemporal segregation of division modes remains a fundamental gap in bacterial cell biology.

*Streptomyces* employ spatiotemporally coordinated regulatory hierarchies to synchronize cell division during developmental transitions (6). The initiation of sporulation is regulated by a family of regulators called “*whi*”(6, 7). The two dynamin-like proteins (DLPs), DynA and DynB, are regulated by WhiH and have been characterized as sporulation-specific cell division proteins contributing to the stabilization of Z-rings in *S. venezuelae* (8). DLPs are a family of large GTPases involved in the remodeling of cellular membranes in eukaryotic cells and are ubiquitous in filamentous actinomycetes and cyanobacteria (9, 10). *In vitro* analysis have shown that bacterial DLPs can form helical filaments and are able to transform membranes into tubular-like structures, although how these proteins function *in vivo* remains enigmatic (11–13).

In this study, we investigated the functional relationship between DynAB and their relation with the *Streptomyces* cell division proteins SepX (14) and SsgB (15) through genetic and microscopic analyses. We revealed that DynAB function as a bifunctional switch and that DynAB activities are modulated by interaction with the developmentally regulated protein SsgB and by the availability of GTP, enabling the segregation of vegetative cross-wall formation from sporulation septation. These findings highlight dynamin-like proteins as key regulators linking cell division to development in multicellular bacteria and provide insights into the evolution and developmental processes of multicellular bacteria.

## Results

### *dynAB* deletion leads to formation of excess cross-walls and to cross-wall-like sporulation septa

In *Streptomyces* sporulation, early sporogenic hyphae are devoid of septa (Fig. 1A-2, B), and then ladders of uniformly-spaced sporulation septa (referred to as septa hereafter) are synchronously formed (Fig. 1A-3) (7, 16). However, septa were arranged irregularly in our *S. coelicolor mtrA* deletion mutant Δ*mtrA* (Fig. S1). Although MtrA regulates multiple development-associated genes (17), it was not known whether MtrA regulates any cell division genes. Therefore, using real-time PCR assays, we compared expression of known cell division genes in Δ*mtrA* and the parental strain M145. *dynA* and *dynB* expression was minimal in Δ*mtrA* (Fig. 1C), suggesting that reduced expression of DynAB may contribute to the impaired sporulation of Δ*mtrA*. DynA and DynB were characterized as necessary for stabilizing Z-rings during sporulation-specific cell division in *S. venezuelae* (8). However, using a Δ*dynAB* mutant, we demonstrated that DynAB impact the formation of division septum and Z-ring in both vegetative hyphae and sporogenic hyphae in *S. coelicolor* M145 (Fig.1D, E, and Fig. S2), indicating the involvement of DynAB in vegetative-specific cell division as well. The average length of the Δ*dynAB* vegetative hyphal compartment was 4.18 µm, nearly half the length of that in M145 (7.95 µm) (Fig. 1D, F), indicating higher frequency of cross-walls in Δ*dynAB* and suggesting that DynAB modulate cross-wall formation in vegetative hyphae. Additionally, the average distance between two adjacent sporulation septa (i.e., spore length) of Δ*dynAB* was 3.19 µm, much longer than in M145 (1.18 µm) and almost equivalent to the interval of Δ*dynAB* cross-walls (Fig. 1E, F), indicating that deletion of *dynAB* results in formation of cross-wall-like septa in sporulation hypha. Collectively, our data suggest DynAB modulate formation of cross-walls in vegetative hyphae and cross-wall-like septa in sporogenic hyphae in *S. coelicolor*.

**Fig. 1.**
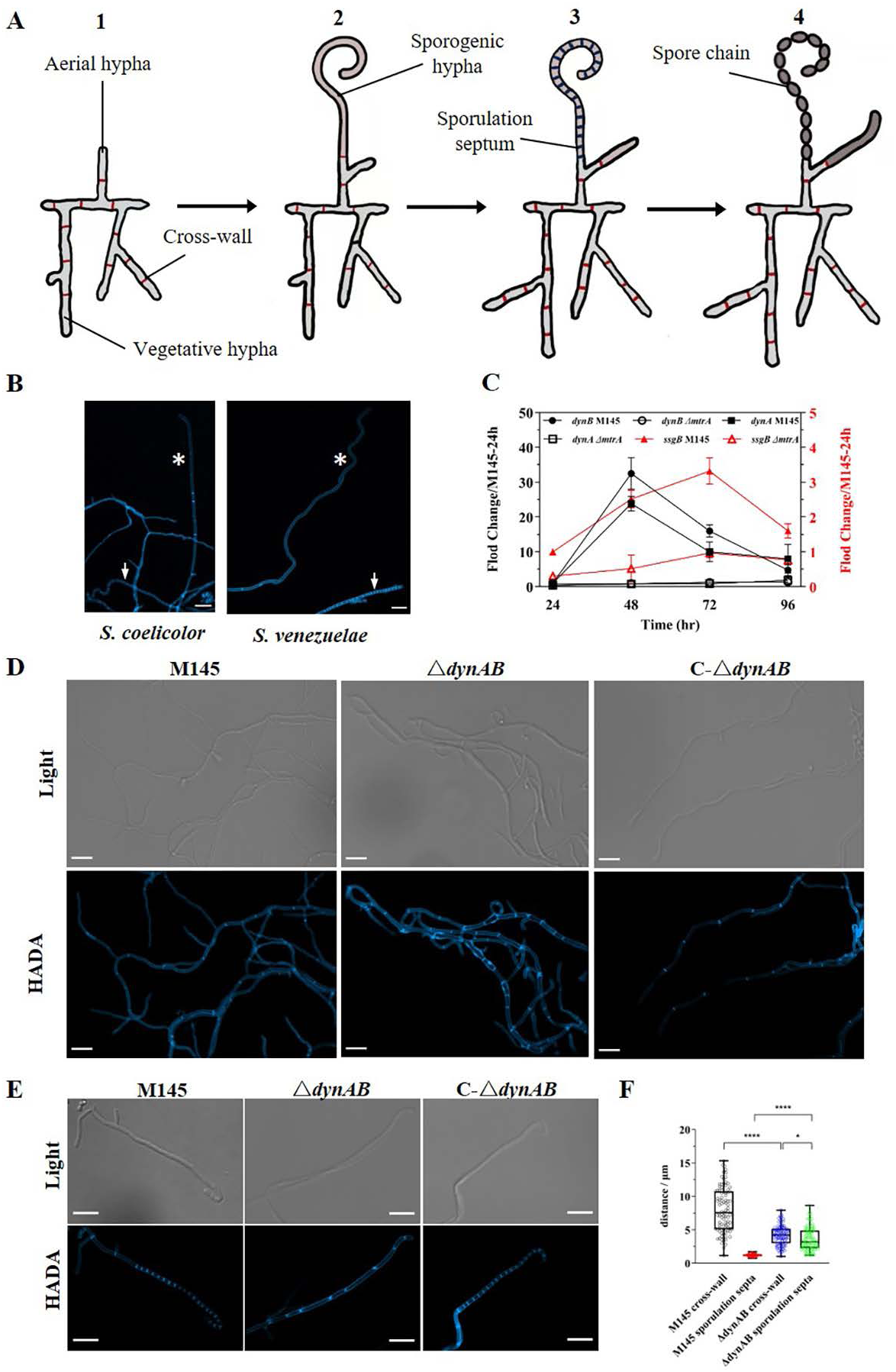
DynAB is required for both vegetative-specific and sporulation-specific cell division in *S. coelicolor*. (A) Schematic representation showing cell division in the development of *Streptomyces* species, which form vegetative hyphae with cross-walls (red bars) and aerial hyphae (A-1); the latter then develop into sporogenic hyphae without sporulation septa at the early stage (A-2), followed by simultaneous formation of sporulation septa (black bars) at the late stage (A-3), which eventually form into chains of spores (A-4). (B) Fluorescence micrographs of early sporogenic hyphae from *S. coelicolor* and *S. venezuelae*. The asterisks indicate early sporogenic hyphae free of sporulation septa, and white arrows indicate ladder-like sporulation septa in late sporogenic hyphae. (C) Temporal expression patterns of *dynAB* and *ssgB* in the *S. coelicolor* wild-type strain M145 and Δ*mtrA*. The parental strain M145 (WT) and Δ*mtrA* were grown on solid BSCA medium, and RNA samples were isolated at the indicated times. Gene expression was measured by real-time PCR, and expression of *hrdB,* encoding the major sigma factor in *Streptomyces*, was used as an internal control. The y-axis shows the fold change in expression level of WT and Δ*mtrA* at each time point over the expression level of each gene in WT at 24 h, which was arbitrarily set to one. Results are the means (±SD) of triplet biological experiments. (D, E) *S. coelicolor* wild-type strain M145, Δ*dynAB*, and the complemented strain C-Δ*dynAB* were grown on BSCA agar supplemented with HADA. (D) Comparison of vegetative hyphae. Light images show the vegetative hyphae, and fluorescence images show the frequent cross-walls formed in the vegetative hyphae of Δ*dynAB* compared to those formed in M145 and the complemented strain C-Δ*dynAB*. Scale bar, 5 μm. (E) Comparison of sporogenic hyphae. Light images show the sporogenic hyphae, and fluorescence images show the ladder-like sporulation septa in M145 and C-Δ*dynAB* and the cross-wall-like sporulation septa in Δ*dynAB*. Scale bar, 5 μm. (F) Hyphal compartment lengths. Statistical analysis of the average length of compartments between two neighboring cross-walls in vegetative hyphae or between septa (or cross-wall-like septa) in sporogenic hyphae in M145 and Δ*dynAB*. Student’s t-test was used for comparison, *ρ<0.05, **ρ<0.01, ****ρ<0.005.

**Fig. 2.**
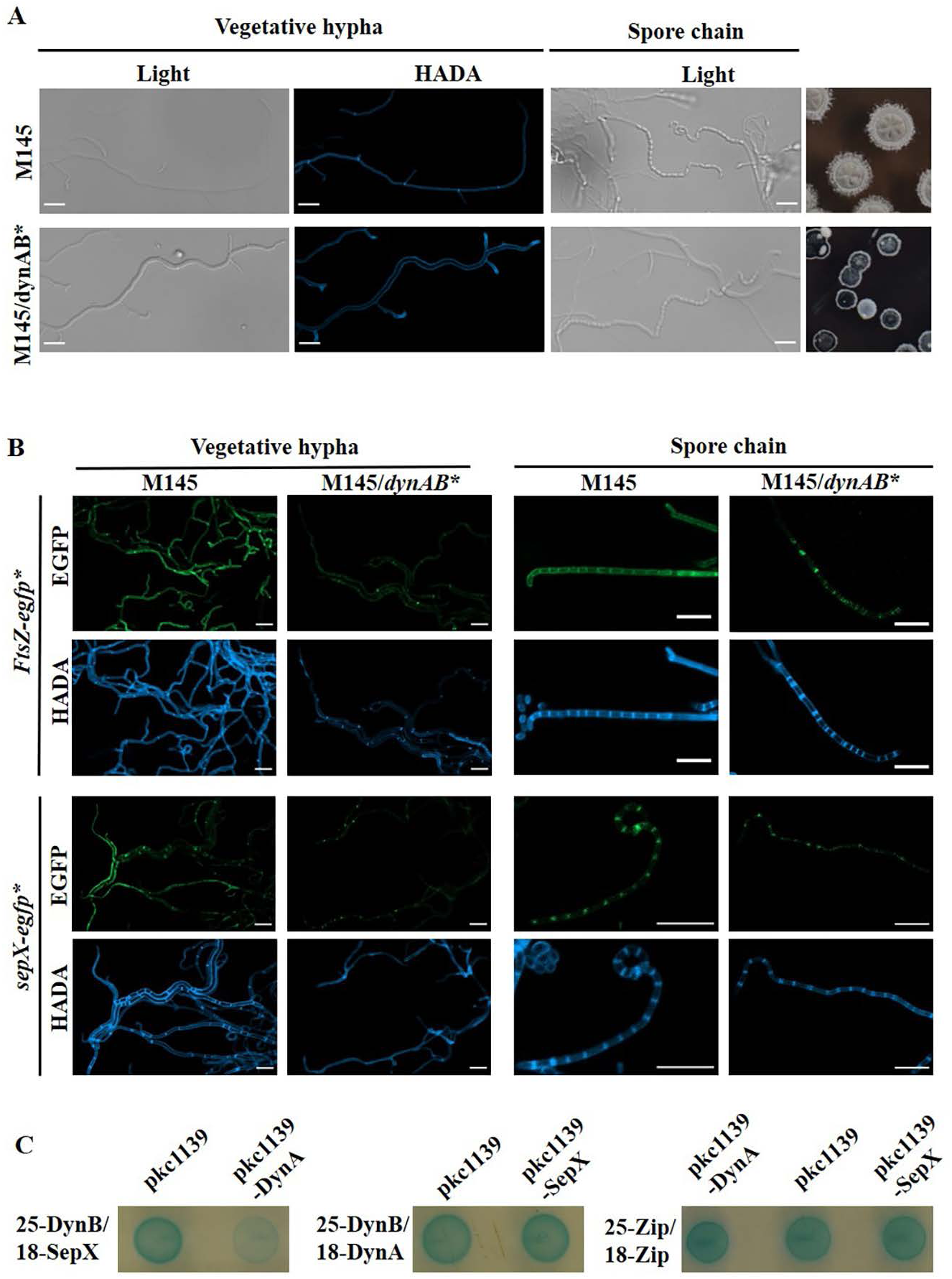
Impact of DynAB on cell division in *S. coelicolor*. (A, B) Wild-type strain M145 and its derivative strains were grown on HADA-supplemented BSCA agar. (A) Effect of constitutive DynAB expression on formation of cross-walls and spore chains in *S. coelicolor*. *dynAB* were expressed from the *kasO* promoter in M145/*dynAB*\*. Fluorescence images show hyphae devoid of cross-walls in M145/*dynAB*\*, and light images show vegetative hyphae and spore chains of both strains. Scale bar, 5 μm. (B) Impact of DynAB on FtsZ and SepX. Gene fusions *ftsZ-egfp* and *sepX*-egfp were expressed from the *kasO* promoter, generating proteins with C-terminal EGFP tags. Green fluorescence images show the localization of the EGFP-tagged proteins in corresponding strains in vegetative or sporogenic hyphae. Blue fluorescence images show the location of cross-walls and sporulation septa in vegetative and sporogenic hyphae, respectively. Scale bar, 5 μm. (C) Bacterial two-hybrid interaction assays with DynA, DynB, and SepX of *S. coelicolor*. Interaction between two fusion proteins generated using pKT25 and pUT18C was evaluated in the presence of a third protein construct based on pKC1139. The blue color indicates protein-protein interactions between the two fusion proteins with stronger color indicating stronger interaction. Interaction between the Zip protein controls was not affected by DynA and SepX, as Zip does not interact with DynA or SepX.

**Fig. 3.**
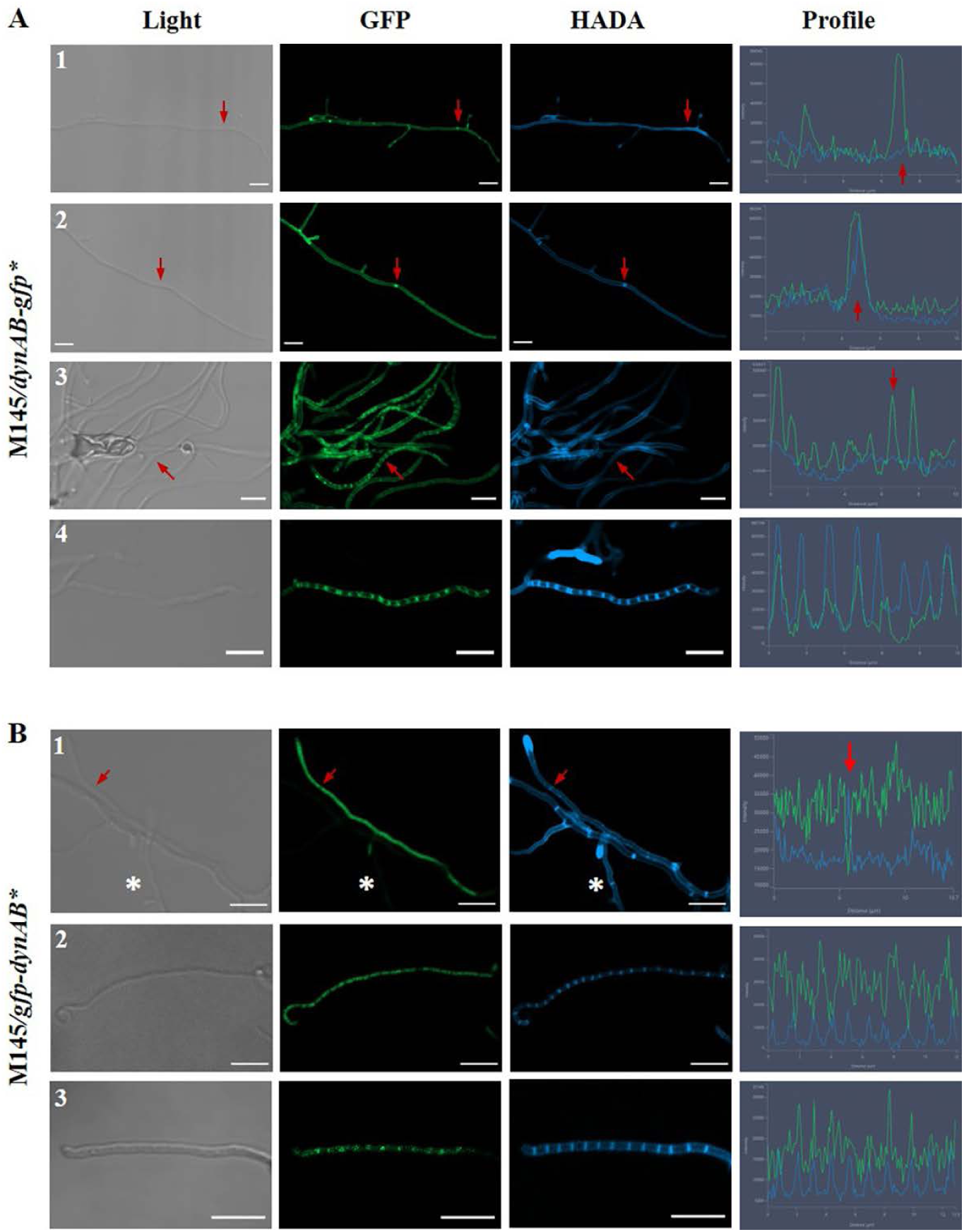
Dynamic localization of DynA and DynB in *S. coelicolor*. (A) DynB localization. Strain M145/*dynAB*-*gfp*\* expresses DynB with a C-terminal GFP-tag, using the gene fusion construct *dynAB-gfp*. M145/*dynAB*-*gfp*\* was grown on HADA-supplemented BSCA agar. Light, green, and blue images show the hyphae, localization of GFP-tagged DynB, and cross-wall or sporulation septa, respectively, of the same view, and demonstrate that DynB is an early cell division protein. DynB co-localized with cross-walls (A-2) and sporulation septa (A-4), with positioning occurring earlier than the formation of cross-walls in vegetative hypha (A-1) or sporulation septa in sporogenic hyphae (A-3). Green and blue electropherograms in the rightmost panels show the strength of the fluorescence emitted by the GFP-tagged DynB and the HADA-stained cross-walls or sporulation septa. Scale bar, 5 μm. (B) DynA localization. Strain M145/*gfp*-*dynAB*\* expresses DynA with an N-terminal GFP-tag, using the gene fusion construct *gfp*-*dynAB*. Additional details as in panel (A) legend. Images demonstrate a highly dynamic localization pattern of DynA with almost continuous distribution, except at cross-wall positions, in vegetative hypha (B-1), whereas DynA alternated with sporulation septa at the early stage in sporogenic hyphae (B-2), and co-localized with sporulation septa at the late stage in sporogenic hyphae (B-3). Rightmost panels, strength of the fluorescence emitted by the GFP-tagged DynA and HADA-stained cross-walls or sporulation septa. Scale bar, 5 μm.

Complementation of Δ*dynAB* with *dynAB* from either *S. coelicolor* or *S. venezuelae* resulted in a phenotype comparable to that of M145 (Fig. 1D, E, and Fig. S3), suggesting a conserved role for DynAB in *Streptomyces*. However, neither *dynA* nor *dynB* alone could complement the defects of Δ*dynAB* and restore normal sporulation (Fig. S3), indicating that both proteins are required. Δ*dynA*, Δ*dynB*, and Δ*dynAB* all formed irregularly-sized spores (Fig. S4), consistent with observations on similar mutants in *S. venezuelae* (8).

### DynAB potentially inhibit SepX function

To verify the inhibitory role of DynAB on cross-wall formation, *S. coelicolor dynAB* were expressed under the constitutive promoter *kasOp*\* (18) in M145. The vegetative hyphae of the resulting strain, M145/*dynAB*\*, were completely devoid of cross-walls (Fig. 2A). Constitutive expression of *S. venezuelae dynAB* in M145, but not *dynA* or *dynB* alone, similarly suppressed cross-wall formation (Fig. S5), indicating that DynA and DynB cooperation is required for inhibiting cross-walls. Although the development of M145/*dynAB*\* was delayed (Fig. S6), spore chains/sporogenic hyphae were much longer than in M145 (Fig. 2A, 4A), suggesting DynAB may promote the growth of early sporogenic hyphae (Fig. 1A-2, 1B) by restricting septum formation.

To test the impact of DynAB on formation of cross-wall-like septa, *dynAB* were constitutively expressed in an *ssgB* mutant, which, like Δ*dynAB*, formed cross-wall-like septa in sporogenic hyphae (Fig. S7). Formation of cross-walls in vegetative hyphae and cross-wall-like septa in sporogenic hyphae was completely inhibited in Δ*ssgB/dynAB\** (Fig. S7), supporting a role for DynAB in repressing these structures. As peak *dynAB* expression overlapped the development of early sporogenic hyphae (Fig. 1A-2, B, C) and abundant DynAB repressed formation of cross-wall-like septa, DynAB appears to play a critical role in maintaining early sporogenic hyphae in a septum-free status, thus facilitating elongation of early sporogenic hyphae.

FtsZ and SepX are the only two *Streptomyces* cell division proteins known to be essential for cross-wall formation (14, 19), as supported by our mutants (Fig. S8). We therefore evaluated the impact of DynAB on SepX and FtsZ localization using the EGFP fusion proteins SepX-EGFP and FtsZ-EGFP. Both SepX-EGFP and FtsZ-EGFP co-localized with cross-walls and septa in M145 (Fig. 2B). FtsZ-EGFP expression did not alter the frequency of cross-walls or septa in M145, whereas its expression restored a more normal number of cross-walls to M145/*dynAB*\* (Fig. 2B); Z-ring formation was visibly impaired in M145/*dynAB*\* (Fig. 2B), indicating that Z-rings are not stable in M145/*dynAB*\*. SepX-EGFP expression appeared to increase the number of cross-walls in M145, consistent with a previous report (14), but had no detectable impact on sporulation-specific division (Fig. 2B). However, the strain co-expressing *dynAB* and *sepX-egfp* formed vegetative hyphae devoid of cross-walls, as with M145/*dynAB*\*, suggesting an inhibitory effect of DynAB on SepX in cross-wall formation. Additionally, SepX-EGFP failed to assemble into ring-like structures in sporogenic hyphae of M145/*dynAB*\* (Fig. 2B), instead forming relatively diffuse patches, suggesting DynAB can repress or prevent the proper assembly of SepX in sporogenic hyphae. Notably, our two-hybrid data indicated that DynA inhibited the interaction between SepX and DynB (Fig. 2C). As described below, DynB is recruited at the early stage of septum formation, and therefore the complexing of DynB with DynA may prevent the recruitment and assembly of SepX by DynB. As SepX is essential for cross-wall formation and for Z-ring formation during sporulation (14), DynAB may control cell division by impacting the recruitment/assembly of SepX.

### Dynamic localization of DynAB

As reported by others (8), DynAB are required for normal sporulation-specific cell division (Fig. 1E and Fig. S4), and our data show that they also repress vegetative-specific cell division (Fig. 1D, 2A), raising the question of how DynAB coordinate these two seemingly opposite functions. Since dynamic localization regulates the activities of DynAB homologues (20, 21), we investigated DynAB localization using strains expressing either EGFP-tagged DynA (M145/*gfp*-*dynAB*\*) or DynB (M145/*dynAB*-*gfp*\*) (Fig. 3). DynB localized to the membrane in vegetative and sporogenic hyphae (Fig. 3A). In young vegetative hyphae, before visible cross-wall formation, DynB was already assembled into tiny foci (Fig. 3A-1) and then co-localized with cross-walls after these emerged (Fig. 3A-2). In early sporogenic hyphae still devoid of sporulation septa, DynB was already assembled into a laddered pattern of ring-like structures (Fig. 3A-3), and then the DynB-ring co-localized with sporulation septa after their formation (Fig. 3A-4). These data indicated that DynB is distributed on the cell membrane and then assembles into Z-ring-like structures before cross-walls or sporulation septa form at future division sites. The cytoplasmic protein DynA was distributed nearly ubiquitously, except for notable absence at sites of cross-wall formation (Fig. 3B-1). In early sporogenic hyphae, DynA was evenly distributed in the sections between the sporulation septa (Fig. 3B-2); these DynA patches then condensed into ring-like structures and co-localized with the sporulation septa in late-stage sporogenic hyphae (Fig. 3B-3). This dynamic localization pattern suggests contrasting roles for DynA in the formation of cross-walls and sporulation septa. Our localization data also suggested that only when the cytoplasmic DynA is complexed with membrane-anchored DynB is the repression of septum formation possible. Based on our data, we hypothesize that, after early cell division proteins are recruited to the incipient division site, DynA dissociates from the DynAB complex, leaving DynB to interact with other cell division proteins to promote the formation of division rings. However, when DynA and DynB interaction is required, such as during constriction of sporulation septa, which enables spore separation, DynA is then re-recruited to the division sites to be complexed with DynB, while this step is not required for cross-wall formation.

**Fig. 4.**
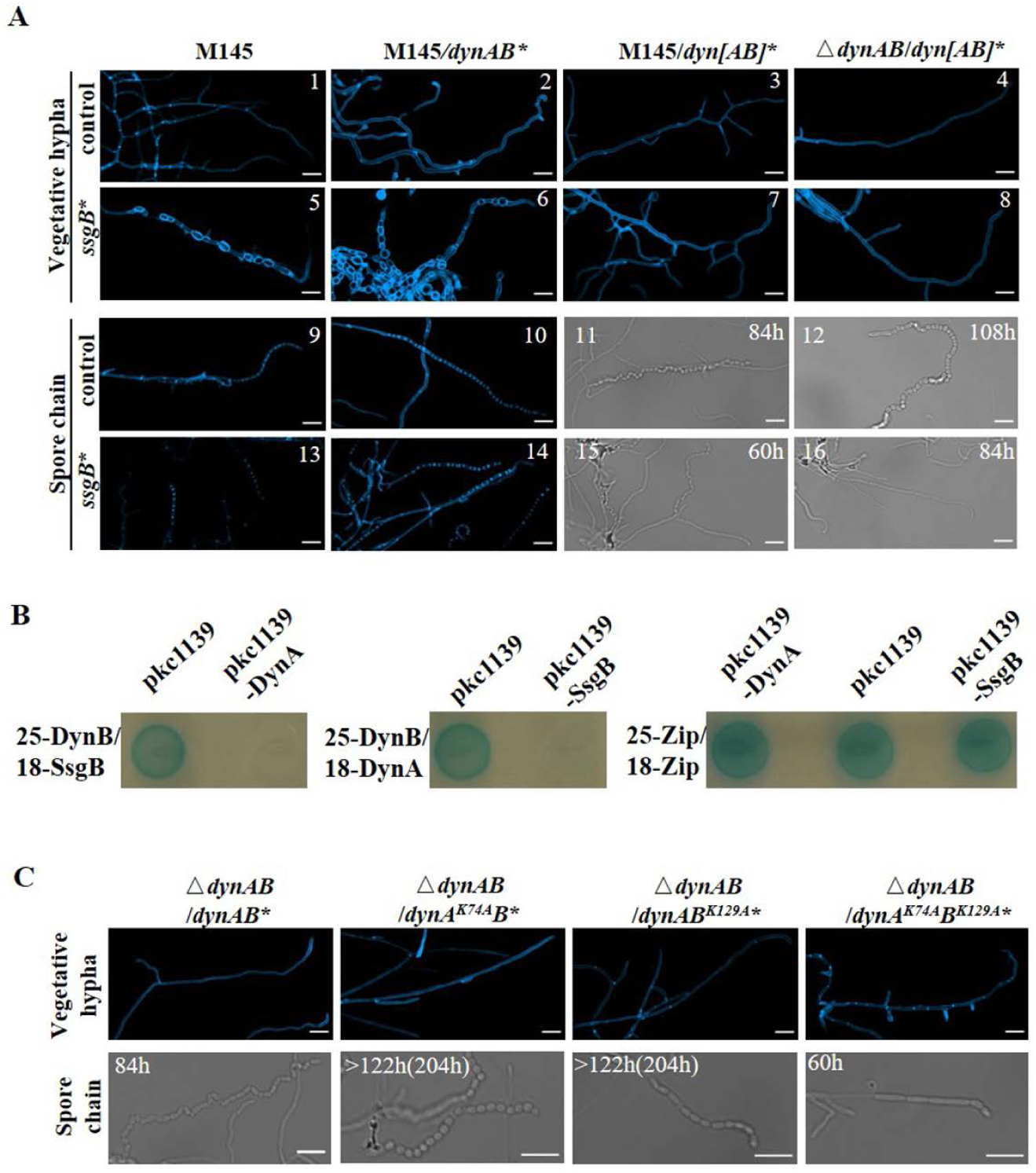
Impact of SsgB and GTP-binding domain mutations on DynAB activity. (A) Separable DynA and DynB are required for SsgB induction of spore-like structures in vegetative hyphae. *dynAB*, *dyn[AB]*, and *ssgB* were expressed from the *kasO* promoter. M145 and derivative strains were grown on HADA-supplemented BSCA agar. Blue fluorescence images show cross-walls in vegetative hyphae and sporulation septa in aerial hyphae. Septum-like cross-walls were detected in M145 expressing SsgB alone or co-expressing SsgB and DynAB, but not with co-expression of SsgB and fused DynAB. Scale bar, 5 μm. (B) Bacterial two-hybrid interaction assays with DynA, DynB, and SsgB of *S. coelicolor*. For details, see Figure 2 legend. Interaction between the Zip protein controls was not affected by DynA and SsgB, which do not interact with Zip. (C) Functional analysis of the GTP-binding domains of DynAB. Mutated proteins DynA(K74A) and DynB(K129A) were expressed individually or together in Δ*dynAB*. Blue fluorescence images show cross-walls in vegetative hyphae, and light images show spore chains in the corresponding strain. Strains with a single GTP-binding domain mutation did not form cross-walls but formed regular spores, whereas the double-mutation strain formed cross-walls and cross-wall-like septa with irregular spores, as in Δ*dynAB* (Fig. 1E). Scale bar: 5 μm.

### Disruption of the DynAB complex by SsgB initiates sporulation-specific cell division

Since DynA was not complexed with DynB in early sporogenic hyphae (Fig. 3B-2), although DynA can strongly interact with DynB (Fig. S9) (8), we speculated that another early cell division protein may disrupt or inhibit formation of the DynAB complex, potentially SepX or SsgB as these proteins also interact with DynB (Fig. S9) (8, 14). SepX could not restore cross-wall formation when co-expressed with DynAB (Fig. 2B), suggesting that SepX is recruited to the division site after the separation of DynA and DynB. In *Streptomyces*, SsgB recruits FtsZ at the initiation of sporulation (15); to test the effect of SsgB on the DynAB complex, *ssgB* was constitutively expressed in M145 and M145/*dynAB*\*, yielding strains M145/*ssgB*\* and M145/*dynAB*\*/*ssgB*\*. Expression of *ssgB* in M145 led to formation of short hyphal compartments in vegetative hyphae (Fig. 4A-5), which were very similar to the sporogenic hyphal compartments of M145 (Fig. 1A-4, Fig. 2A).

**Fig. 5.**
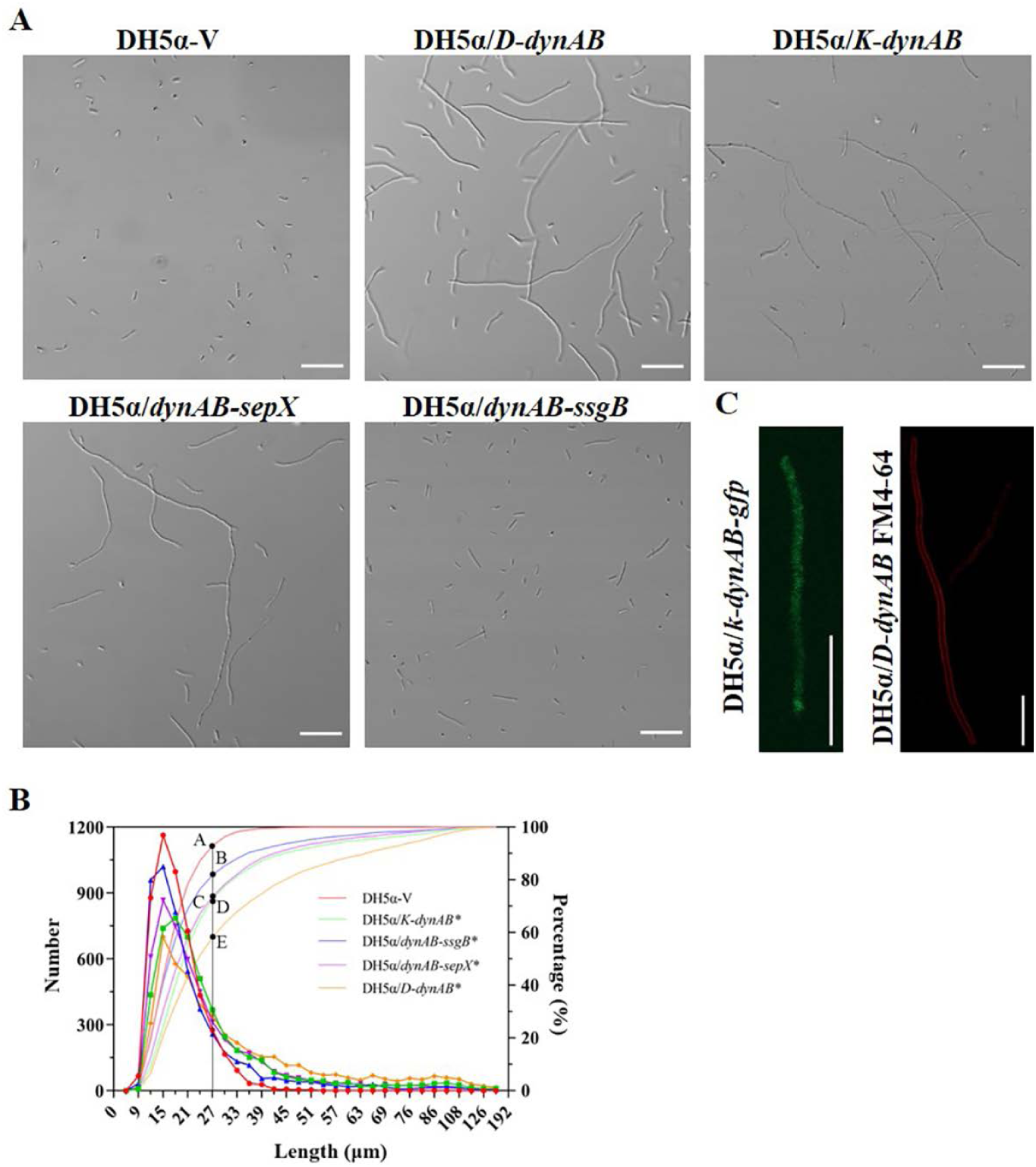
Formation of unicellular, filamentous cells by *E. coli* expressing *Streptomyces dynAB.* (A) Microscopic images of cell cultures from derivatives of *E. coli* strain DH5α. DH5α/K-*dynAB*\* and DH5α/D-*dynAB*\* express *dynAB* from, respectively, the *kasOp\** promoter and the *rrnD* promoter; DH5α/*dynAB*-*sepX*\* and DH5α/*dynAB*-*ssgB*\* co-express *sepX* and *ssgB*, respectively, with *dynAB*. DH5α/V, control strain containing the cloning vector. Higher numbers of long filamentous cells were detected in DH5α/K-*dynAB*\*, DH5α/D-*dynAB*\*, and DH5α/*dynAB*-*sepX*\*. (B) Flow cytometry analysis showing the distribution and percentage of cells with different lengths for *E. coli* strains shown in panel (A). (C) Fluorescence image showing uniform distribution of DynB in DH5α/K-*dynAB-gfp* and absence of cell division septa in long filamentous cell of DH5α/K-*dynAB* visualized by FM4-64 staining. Scale bar, 20 μm.

M145/*dynAB*\*/*ssgB*\* also formed short hyphal, spore-like compartments (Fig. 4A-6), like those of M145/*ssgB*\*; therefore, the cross-wall-free vegetative hyphae observed with M145/*dynAB*\* (Fig. 4A-2) became loaded with septum-like cross-walls upon SsgB overexpression, suggesting disruption of DynAB complexes by SsgB. Strain M145/*ssgB*\* also formed short sporogenic hyphae containing about 10 septa, compared to the longer sporogenic hyphae of M145 with about 15 septa, suggesting abundant SsgB reduces growth of early sporogenic hyphae. In contrast, M145/*dynAB*\*/*ssgB*\* formed ladder-like septa of normal length (Fig. 4A-14), implying that DynAB and SsgB are antagonistic during growth of early sporogenic hyphae and initiation of sporulation-specific cell division. We deduce that, by preventing cell division in early sporogenic hyphae, DynAB activity favors formation of long sporogenic hypha, whereas SsgB leads to premature cell division in sporogenic hyphae. The two-hybrid data with DynA, DynB, and SsgB (Fig. 4B) indicate that DynA and SsgB compete for DynB, and our overall data suggest that the outcome of this competition influences the progress of sporogenic hyphal development. According to their temporal patterns (Fig. 1C), *dynAB* expression peaks when early sporogenic hyphae are developing, before peak expression of *ssgB*, thereby enabling *Streptomyces* to control the length of sporogenic hyphae and consequently the number of spores.

**Fig. 6.**
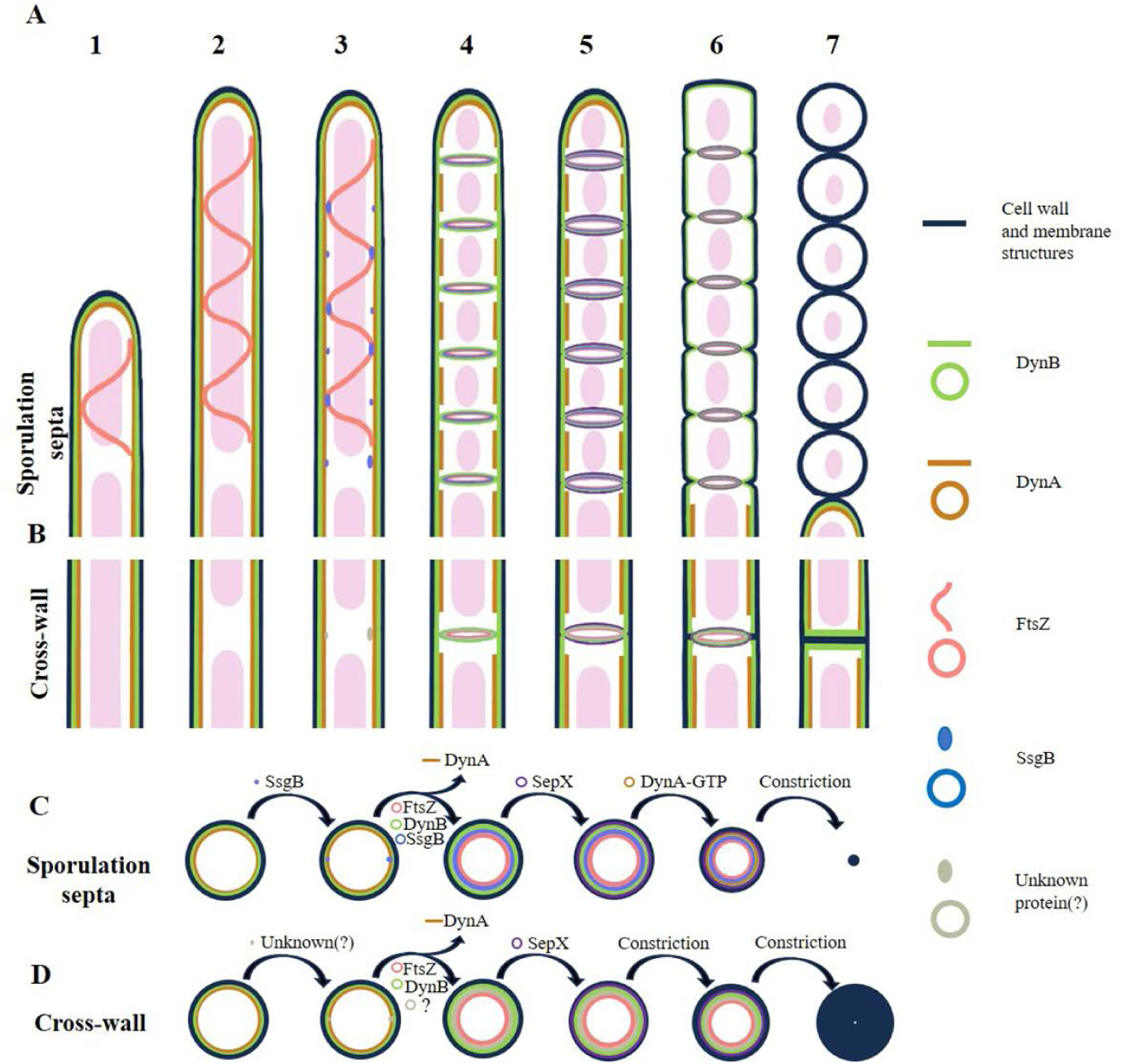
A proposed model for the working mechanism of DynAB in *Streptomyces*. (A, B) Longitudinal section and (C, D) cross-section of (A, C) sporogenic hyphae and (B, D) vegetative hyphae showing formation of sporulation septa and cross-walls. In this model, DynA interacts with the membrane-bound DynB, forming DynAB complexes (steps 1 and 2), which can further polymerize into spiral/tubular filamentous structures embedded in the inner membrane surface, preventing cross-wall formation. However, when SsgB is highly expressed (step 3), the DynAB complexes at division sites dissociate (step 4), allowing DynB to interact with SsgB and then form the early divisome, together with FtsZ (step 4). SepX is then recruited (step 5), working alongside DynB and other cell-division proteins to initiate the formation of sporulation septum. At the initiation of constriction, DynA rebinds DynB of the divisome and constricts together with the septum upon GTP hydrolysis (steps 6 and 7), ultimately separating the sporulation hyphae into dozens of spores.

Like *Streptomyces dynAB*, bacterial DLP genes often exist as tandem pairs (10). However, *Bacillus subtilis dynA* is a fusion of two dynamin genes containing two GTPase domains that function in membrane remodeling (11). To test if the separation of DynAB by SsgB is essential for septum formation and the suppression of cross-wall formation, we generated the DynA and DynB fusion construct Dyn[AB] to prevent DynAB separation. When Dyn[AB] was expressed in M145 (M145/*dyn[AB]*\*), vegetative cross-walls were not formed (Fig. 4A-3), indicating that Dyn[AB] could still repress cross-wall formation. Notably, with Dyn[AB] expression in M145/*ssgB*\* (M145/*ssgB*\*/*dyn[AB]*\*), the short spore-like compartments found with M145/*ssgB*\* were not formed (Fig. 4A-7); therefore, unlike DynAB, Dyn[AB] could repress the excessive septum-like cross-wall formation resulting from SsgB overexpression, implying that SsgB must dissociate the DynAB complex in order to induce spore-like compartments in vegetative hyphae. As dissociation of the DynAB complex is also required in forming sporulation septa, Dyn[AB] should impact formation of sporulation septa. Whereas expression of the fused Dyn[AB] did not inhibit spore formation in M145, the length of sporogenic hyphae was extended (Fig. 4A-11), with normal lengths restored when Dyn[AB] was co-expressed with SsgB (Fig. 4A-15), similar to DynAB. We deduced that the failure of Dyn[AB] to inhibit spore formation could be potentially caused by the native DynAB in M145; therefore, Dyn[AB] was expressed in Δ*dynAB*. As expected, formation of cross-walls was inhibited in Δ*dynAB*/*dyn[AB]* (Fig. 4A-4). Furthermore, the cross-wall-like septa observed in Δ*dynAB* were absent; instead, nearly normal spores were formed in Δ*dynAB*/*dyn[AB]* (Fig. 4A-12), although their formation was markedly delayed when compared with Δ*dynAB* spores, suggesting that Dyn[AB] could still repress formation of cross-wall-like septa, and the much-delayed sporulation could be caused by an impaired divisome due to the inseparable Dyn[AB]. Additionally, a Δ*dynAB* strain that co-expressed Dyn[AB] with SsgB was free of vegetative cross-walls but formed spores of irregular lengths (Fig. 4A-16), suggesting the premature formation of spores by SsgB. Collectively, our data suggest that disruption of DynAB complexes by SsgB initiates sporulation-specific cell division in *Streptomyces*.

### DynAB GTP-binding activity impacts release and recruitment of DynA

When sporulation septum formation begins, DynA is not complexed with DynB; however, DynA is eventually recruited to the division site, forming DynAB complexes (Fig. 3B). DynB recruits DynA to the divisome, and this process depends on the GTP-binding ability of both proteins in *S. venezuelae* (8). With paired bacterial DLPs, GTP can regulate the interaction and polymerization of the two subunits (8, 13). To test the potential impact of GTP-binding by DynAB on cell division, essential GTP-binding sites in the DynA or DynB GTP domain P-loop were mutated (8), resulting in DynA^K74A^B, DynAB^K129A^, and DynA^K74A^B^K129A^, which were expressed in Δ*dynAB* (Fig. 4C). Cross-walls were not formed in the single P-loop mutants Δ*dynAB*/DynA^K74A^B* and Δ*dynAB*/DynAB^K129A^*, indicating that DynAB with one mutation could still repress cross-wall formation, suggesting DynAB complexes still formed through other weak interactions between these two proteins (8). However, cross-walls were restored in Δ*dynAB*/DynA^K74A^B^K129A^*, indicating that DynAB with mutations in both loops failed to repress cross-wall formation, likely due to loss of interaction between these two mutated proteins and the resulting absence of functional DynAB complexes.

Notably, DynAB with a single mutation expressed by its native promoter could not restore normal spore formation in Δ*dynAB* (8), whereas constitutively expressed DynAB with a single mutation (DynA^K74A^ or DynB^K129A^) restored normal spore formation in Δ*dynAB* (Fig. 4C), albeit with much-prolonged growth time, similar to the result with Δ*dynAB*/*dyn[AB]*\*. Therefore, DynAB with a single mutation or fused Dyn[AB] could still repress the formation of cross-wall and cross-wall-like septa, but may fail to promote the constriction of septa. Expression of these mutated proteins in M145 resulted in similar phenomena in cross-wall formation (Fig. S10). These findings suggested that GTP-binding ability modulates the interaction between DynA and DynB and thereby the physiological function of these two proteins. Therefore, the effects of GTP levels were evaluated using the GTP-sensing protein GEVAL30 in *Streptomyces* (22). Fluorescence analysis detected higher GTP levels in sporogenic hyphae than in other cells (Fig. S11), which would aid in the recruitment of GTP-bound DynA to the division septum by GTP-bound DynB.

### DynAB effects on cell division in *Escherichia coli*

Although DynAB are conserved in filamentous actinobacteria (8, 10), structurally similar proteins are not found in single-celled bacteria, such as *E. coli* DH5α. Due to their ability to suppress *Streptomyces* cell division, we tested whether *Streptomyces* DynAB could inhibit binary division in *E. coli* DH5α, by generating strains DH5α/D-*dynAB* and DH5α/K-*dynAB,* which express *dynAB* from the *E. coli rrnD* (23) promoter and *kasOp\**, respectively. Microscopic analysis showed that DH5α/D-*dynAB* and DH5α/K-*dynAB* cells were dramatically longer than the control DH5α-V cells (Fig. 5A). Quantification of elongated cells by flow cytometry showed that the majority of DH5α/V cells (92.98%) were shorter than 27 µm (Fig. 5B). Taking this length as a threshold, approximately 7.02% of cells in DH5α/V were longer, potentially due to the antibiotic added in the medium (24). By contrast, elongated cells accounted for 27.41% of DH5α/K-*dynAB* cells and 41.68% of DH5α/D-*dynAB* cells, indicating a profound effect of DynAB on the formation of filamentous cells in *E. coli,* with the *E. coli* promoter *rrnD* having a greater effect than the *Streptomyces* promoter *kasOp\**. Using GFP-tagged constructs, fluorescence microscopy revealed that DynB-GFP was uniformly distributed in DH5α/K-*dynAB-gfp*, and no cell division septum was observed in the elongated, filamentous cells of DH5α/D-*dynAB* when using FM-64 staining (Fig. 5C). Consistent with the long, cross-wall-free vegetative hyphae caused by DynAB overexpression in *Streptomyces*, our data suggest that DynAB can inhibit *E. coli* cell division and transform short, rod-shaped *E. coli* cells into filamentous forms.

According to our data, the ability of DynAB to maintain early sporogenic hyphae in a septum-free state could be relieved by SsgB (Fig. 4A-6), but not SepX (Fig. 2B). Therefore, we tested the impact of SsgB and SepX co-expression with DynAB (under the control of *kasOp*) on formation of filamentous *E. coli* (Fig. 5A). DH5α/*dynAB*-*sepX* formed long cells, similar to DH5α/D-*dynAB* and DH5α/K-*dynAB* cells, whereas long cells were minimally observed in DH5α/*dynAB*-*ssgB* (Fig. 5A). Flow cytometry demonstrated that 26.39% of DH5α/*dynAB*-*sepX* cells were elongated (Fig. 5B), a rate comparable to that of DH5α/K-*dynAB*. However, co-expression of *dynAB* with *ssgB* resulted in only 18.19% elongated cells, markedly lower than observed in DH5α/K-*dynAB*, suggesting that SsgB can inhibit the ability of DynAB to induce formation of filamentous *E. coli*. Collectively, our data indicate that the *Streptomyces* DynAB complex can inhibit cell division in *E. coli* and that SsgB can disrupt these complexes and trigger cell division in *E. coli*, consistent with the roles of DynAB and SsgB in *Streptomyces*.

## Discussion

Although the genus *Streptomyces* has been studied for nearly a century (25)(17)(19), the mechanisms that control vegetative and sporogenic cell division are still poorly understood. In this study, we discovered that the two DLPs DynAB of *Streptomyces* not only function as sporulation-specific cell division proteins, as reported (8), but also as a negative regulator of cell division by repressing cross-wall formation in vegetative hyphae and preventing the formation of aberrant cross-wall-like sporulation septa in sporulation hyphae, thereby favoring elongation of sporogenic hyphae. Importantly, we also discovered that initiation of sporulation-specific cell division requires disruption of DynAB complexes by the FtsZ-recruiting protein SsgB.

We propose two possible mechanisms for the negative impact of DynAB on cell division. One mechanism would be by preventing divisome assembly. In our study, constitutive expression of *dynAB* in vegetative hyphae, a situation that mimics the high level expression of DynAB in sporogenic hyphae, abolished the recruitment or assembly of SepX, which is essential for cross-wall formation (14); SepX recruitment should occur after recruitment of FtsZ, but prior to synthesis of cross-wall peptidoglycan, distinguishing it from the Min system (26, 27), which functions before Z-ring assembly. Another mechanism could be by the assembly (or polymerization) of the DynAB complex itself. DynAB share structural similarities with the bacterial proteins BDLP12 and Cj-DLP12 (Fig. S12-S14), polymerization of which can drive liposomes into tubular structures *in vitro* (12, 13, 21). When assembled *in vivo*, DynAB may create steric barriers that physically exclude membrane constriction, as supported by heterologous expression of DynAB in *E. coli*, which resulted in formation of filamentous cells.

Synchronous sporogenic septation in *Streptomyces* is preceded by the synchronous assembly of a large number of regularly spaced Z-rings along the sporogenic hypha (8, 28). Deletion of *dynAB* disrupted Z-rings in the sporogenic hyphae of M145 (this study) and *S. venezuelae* (8), but DynAB do not appear to aid in determining division sites. Although SsgB is known to recruit FtsZ to sporogenic division sites (15), in this study, we showed that SsgB can also disrupt the interaction between DynAB, thereby initiating sporulation-specific cell division. We found that *dynAB* peak expression occurred well before that of *ssgB*, thereby allowing DynAB complexes to form first, thus inhibiting septum formation in the early sporogenic hyphae and enabling their extension. However, our findings suggest that, when SsgB later reaches threshold levels, SsgB can disrupt these DynAB complexes, alleviating the inhibitory effect of DynAB throughout the sporogenic hyphae and initiating synchronous sporulation-specific cell division; thus, the interaction of DynAB and SsgB would serve as a checkpoint for the synchronous initiation of sporulation-specific cell division.

Using fluorescence microscopy to observe the dynamic localization of DynA, we found that DynA released in free form from DynAB complexes by SsgB can then be recruited by DynB when sporulation septa start constriction. This process may rely on the GTP-binding activities of DynAB (29). In *B. subtilis*, a single GTPase domain loop mutant decreased the self-interaction of DynA (11), and a similar phenomenon was observed between DynA and DynB in *S. venezuelae* (8) and in our study. Although mutation of the GTPase loop in either DynA or DynB decreased the interaction strength and failed to restore normal spore length in *S. venezuelae* (8), we found that constitutively expressed DynAB with a single mutation could still inhibit formation of cross-walls and cross-wall-like septa, although with delayed spore formation. As single-loop mutants of DynA in *B. subtilis* and Cj-DLP12 in *C. jejuni* also failed to hydrolyze GTP (11, 13), the GTP hydrolysis by DynAB in *Streptomyces* may be essential for septum constriction, which would explain why bacteria DLPs do not require GTP to promote formation of tubular structures in vitro, but do need GTP to catalyze membrane cleavage (12, 13, 30).

Structural defects observed in the rare DynAB homologues (e.g., truncated GTPase domains) found in unicellular bacteria further indicate genomic erosion following the loss of an original function, reinforcing the hypothesis that DynAB systems may represent an evolutionary adaptation in sophisticated prokaryotic development.

Based on our findings, we propose a model for the simultaneous formation of sporulation septa in sporogenic hyphae (Fig. 6). Polymerized DynAB complexes attach to the inner membrane surface to keep early sporogenic hyphae septum-free. However, when SsgB is expressed, SsgB interacts with the DynAB complexes, displacing DynA. About the same time, FtsZ is recruited to the future division site by SsgB to form the early divisome. SepX is recruited to the divisome by DynB after separation of DynAB, initiating formation of the sporulation septum. The high GTP level in late sporogenic hyphae allows binding of GTP by DynA, which re-enables DynA and DynB interactions. Consequently, GTP-bound DynA is recruited to the sporulation divisome by DynB; then, under GTP hydrolysis by FtsZ and DynAB, the septum constricts until the spores are separated. In vegetative hyphae, GTP levels are minimal, which does not favor recruitment of DynA or constriction of Z-rings, and therefore sparsely separated cross-wall structures are formed. As SsgB is not required for cross-wall formation, and its level is only minimal during vegetative growth (15), other unknown proteins potentially disrupt DynAB complexes and recruit FtsZ for cell division in vegetative hyphae. Overall, our findings indicate key roles for DynAB in both types of *Streptomyces* cell division through the regulation of cross-walls and sporulation septa and provides insights into how bacteria DLPs may have evolved to have diverse functions in developmentally complex bacteria.

## Materials and Methods

### Bacterial strains and culture conditions

Bacterial strains and plasmids used in this study are listed in Table S1. *S. coelicolor* strain M145 was used as the wild-type strain, and M145 and its derivatives were grown at 30°C on BSCA agar for microscopic analysis and RNA extraction (31), and on ISP3 agar for spore production and conjugal transfer. *Escherichia coli* strains were cultivated in Luria-Bertani (LB) liquid medium. Antibiotics were added, as appropriate, to growth medium for selection of either *E. coli* transformants or *Streptomyces* conjugants.

### Deletion of genes from *S. coelicolor* M145

A CRISPR-Cas9 approach was used to delete genes from M145 as reported (32, 33). The general procedure is described here, using the deletion of *dynA* from M145 as an example. Three primer pairs, *dynA*-sgRNA-F/R, *dynA* Up-F/R, and *dynA* Down-F/R, were designed to generate overlapping sequences between amplified fragments and were used to amplify the guide RNA (sgRNA) for *dynA*, and the upstream and downstream homologous arms of the desired deleted sequence of *dynA*, respectively. These three PCR fragments were purified, mixed, and ligated with vector pKCcase9d6424 (32), pretreated with *Bcu*I and *Hind*III, to generate pKC-*dynA*, which was then transformed into *E. coli* ET12567 (pUZ8002). Conjugation was performed using M145 and *E. coli* ET12567 (pUZ8002) containing pKC-*dynA*. Conjugants resistant to apramycin were selected at 30°C and were then further incubated at 37°C for several rounds to cure the temperature-sensitive vector. Mutants with double-crossovers (Δ*dynA*) were verified by PCR analysis and sequencing. Mutant strains with knockouts of *dynB* (Δ*dynB*), *dynAB* (Δ*dynAB*), *ftsZ* (Δ*ftsZ*), *sepX* (Δ*sepX*), and *ssgB* (Δ*ssgB*) were obtained similarly. Primers used in the generation of constructs used in this study are listed in Table S2.

### Complementation of Δ*dynAB*

The primer pair *dynAB*-Com-F/R was used to amplify a fragment containing *dynAB* and its upstream intergenic sequence in *S. coelicolor*. The PCR product was cloned into the *Hind*III-digested integrative vector pMS82(34) to obtain pCOM-*dynAB*, which was then introduced into *E. coli* ET12567 (pUZ8002) after sequencing. Conjugation was performed using Δ*dynAB* and ET12567 (pUZ8002) containing pCOM-*dynAB*, and the complemented strain C-Δ*dynAB* was obtained after selection by hygromycin and PCR verification. Similarly, C_VEN_-Δ*dynAB*, another complementation strain of Δ*dynAB*, was constructed using *dynAB* and its native promoter from *S. venezuelae*.

### Construction of *S. coelicolor* strains expressing GFP-tagged proteins

Two gene fusions, *kasOp*-*gfp*-*dynAB* and *kasOp*-*dynAB*-*gfp*, which express GFP-tagged DynA and GFP-tagged DynB, respectively, were constructed to determine the subcellular localization of DynA and DynB in M145. *Streptomyces* strains expressing other proteins tagged with GFP either at the N or C terminal were constructed with a similar strategy. Here, we describe the construction procedure for obtaining the M145 strain expressing fusion protein FtsZ-GFP. In this study, most protein genes were expressed from the *kasOp*, a constitutive promoter optimized for gene expression in *Streptomyces* hosts(18). Firstly, primer pair KasOp-F/-R was used to amplify the approximately 100 bp *kasO* promoter sequence using a synthetic gene containing the *kasO* promoter as template (18). Then, a 1.2 kb fragment containing the *ftsZ* coding sequence was amplified using primer pair FtsZ-F/-R and M145 genomic DNA as template. Next, a 450 bp fragment containing the coding sequence of *egfp* was amplified using primer pair EGFP-F/-R and the commercial vector pEGFP-C1 as template. Overlapping sequences in the primers enabled these PCR fragments to be ligated in the order of *kasOp*-*ftsZ*-*egfp* using the ClonExpress®Ultra One Step Cloning Kit (Vazyme), following by cloning into pMD-18T. After sequencing verification, the 1.75 kb insert containing *kasOp*-*ftsZ*-*egfp* was excised by *Kpn*I and *Hind*III digestion and then ligated into the integrative vector pMS82 (34), pretreated with the same two enzymes, to obtain plasmid p*ftsZ*-*gfp*, which was transformed into *E. coli* ET12567 (pUZ8002). Conjugation was performed using M145 and *E. coli* ET12567 (pUZ8002) containing p*ftsZ*-*gfp*. Conjugants resistant to hygromycin were selected at 30°C and were verified by PCR analysis, obtaining M145/*ftsZ*-*gfp*\*, the M145 strain expressing GFP-tagged FtsZ protein. A similar approach was followed for constructing derivative M145 strains expressing GFP-tagged DynA, DynB, SepX, and SsgB proteins. For additional GFP constructs, see the next section.

### Construction of *Streptomyces* strains expressing *Streptomyces* genes

Protein-coding genes were expressed from the *kasO* promoter (18). For constructing strains expressing *dynAB*, primer pair KasOp-F/-R was used to amplify the *kasO* promoter. Then, a 4.09 kb fragment containing the *dynAB* coding sequence was amplified using primer pair DynAB-F/-R and M145 genomic DNA. Overlapping sequences in these primers enabled these two PCR fragments to be ligated in the order of *kasOp*-*dynAB*, using the ClonExpress®Ultra One Step Cloning Kit, followed by cloning into pMD-18T. After sequencing verification, the 4.1 kb insert containing *kasOp*-*dynAB* was excised by *Kpn*I and *Hind*III digestion and then ligated with *Kpn*I/*Hind*III-digested pMS82 (34) to obtain plasmid pST-DynAB, which was transformed into *E. coli* ET12567 (pUZ8002). Conjugants were generated and verified as described for the GFP-tagged constructs, obtaining M145/*dynAB*\*, the M145 strain that constitutively expresses DynAB protein. Derivative M145 strains expressing *dynA*, *dynB*, *ftsZ*, *sepX*, or *ssgB* were similarly obtained.

To generate strain M145/*dynAB*-*ftsZ*\*, which co-expresses *dynAB* and *ftsZ*, gene fusion *kasOp*-*ftsZ* was prepared as described above and inserted into pST-DynAB, obtaining pST-DynAB-FtsZ, which was used for conjugation as described above. M145 strains co-expressing *dynAB* and *sepX* or *dynAB* and *ssgB* were constructed similarly.

For localization of FtsZ in M145/*dynAB*\*, gene fusion *kasOp*-*ftsZ-gfp* was inserted into pST-DynAB, obtaining pST-DynAB-FtsZ-GFP, which was used for conjugation, generating M145/*dynAB*-*ftsZ-gfp*\*, the M145/*dynAB*\* strain expressing GFP-tagged FtsZ protein. M145/*dynAB*\* strains expressing GFP-tagged SepX or SsgB protein were similarly obtained.

### Construction of *E. coli* strains expressing *Streptomyces* genes

To express *dynAB* in *E. coli*, the *kasOp*-*dynAB* construct, described above, was cloned into pCE-Zero, obtaining pKas-DynAB. After sequencing verification, pKas-DynAB was transformed into DH5α, and transformants were selected by antibiotic and verified by PCR analysis, obtaining DH5α/*kasOp*-*dynAB*\*. DH5α expressing *dynAB* from the *rrnD* promoter of *E. coli* was constructed similarly.

To co-express *dynAB* and *sepX* in *E. coli*, gene *sepX* was placed under the control of the *amp* promoter and inserted into pKas-DynAB, obtaining pKas-DynAB-SepX. After sequencing verification, pKas-DynAB-SepX was transformed into DH5α, and transformants were selected using antibiotics and verified by PCR analysis, obtaining DH5α/*dynAB-sepX*\*. The DH5α strain co-expressing *dynAB* and *ssgB* was obtained in a similar way.

### Construction of *Streptomyces* strains expressing DynAB protein mutants

Firstly, primer pair KasOp-F/DynA-MR (carrying the desired mutated nucleotide sequence) and template *kasOp*-*dynAB* constructed as above were used to amplify a 322-bp fragment containing the *kasO* promoter sequence and partial *dynA* coding sequence, and then a 3278-bp fragment containing partial *dynA* (with mutation) and *dynB* was amplified using primer DynA-MF (complementary to DynA-MR) and primer R2 and *kasOp*-*dynAB*. Next, these two fragments were ligated with pretreated pMS-82, obtaining pK-*dynA*(*K74A*)*B*, which was then introduced into M145 and its derivatives through conjugation, obtaining M145/*dynA*(*K74A*)*B*\*; similarly, strains M145/*dynAB*(*K129A*)* and M145/*dynA*(*K74A*)*B*(*K129A*)* were obtained. All these constructs were verified by sequencing.

For constructing strains expressing fused DynAB, we used *kasOp*-*dynAB* as template with primer pair KasOp-F/DynAB-FR, which linked *dynA* and *dynB* without the stop codon of *dynA*, the intergenic sequence, or the start codon of *dynB*, to amplify a 1735 bp fragment; then a 1857 bp fragment was amplified using primers DynAB-FF (complementary to DynAB-FR) and primer R2. Next, these two fragments were ligated with pretreated pMS-82, obtaining pK-*dyn[AB]*, which was then introduced into M145 and its derivatives through conjugation, obtaining M145/*dyn[AB]*\*. All these constructs were verified by sequencing.

### RNA isolation, reverse transcription-PCR (RT-PCR), and real-time PCR

To extract RNA, *S. coelicolor* strains were grown at 30°C on solid BSCA medium covered with plastic cellophane, and the mycelia were collected at various times as indicated. Crude RNA samples were treated twice with ‘Turbo DNA-free’ DNase reagents (Invitrogen) to remove chromosomal DNA. Reverse transcription and real-time PCR assays were carried out as described (17). The SYBR Premix Ex Taq (TaKaRa) was used under recommended conditions on a Roche LightCycler480 thermal cycler to determine the melting curve of PCR products and their specificity. Relative quantities of cDNA were normalized for the *hrdB* gene, encoding the major sigma factor of *Streptomyces*.

### Bacterial two-hybrid interaction assays

The interaction assays were performed essentially as described (35). Briefly, *S. coelicolor* genes were amplified using primers that carried either an *Xba*I or a *Cla*I site (Table S2). The PCR fragments were then purified and inserted into plasmids pKT25 and pUT18 to generate translational fusions with the catalytic domains of *Bordetella pertussis* adenylate cyclase (36, 37). The compatible translational fusion constructs were co-transformed into the *E. coli cya* mutant BTH101, and the resulting transformants were first induced in LB liquid culture containing isopropyl β-D-1-thiogalactopyranoside (IPTG) for protein expression and then spotted on LB plates containing IPTG and 5-bromo-4-chloro-3-indolyl-β-D-galactopyranoside, and incubated at 30°C to detect protein interactions. All constructs were verified by sequencing.

### Microscopic analysis

Spores of *S. coelicolor* strains were plated onto BSCA agar without or with (for fluorescence imaging) addition of HADA (7-hydroxycoumarin 3-carboxylic acid-amino-D-alanine), and sterile coverslips were inserted at an angle to allow growth of *Streptomyces* cells onto the coverslip surfaces. After cultivation at 30°C for the desired time, the coverslips were removed and sealed with oil for microscopic observation. All images were acquired using a Zeiss LSM900 laser scanning confocal microscope.

### Flow cytometry analysis

To determine the numbers of cells of various lengths, an individual clone of each *E. coli* strain was inoculated into LB broth, cultured overnight at 37°C with rotation at 800 rpm, and then inoculated as seed (at 2% volume) into fresh LB broth and incubated until the culture reached an OD_600nm_ value of about 0.5. To avoid potential cell aggregation caused by long filamentous cells, 200 μl of *E. coli* culture was directly mixed with 500 μl PBS buffer (1×), and the mixture was subjected to flow cytometry analysis using the Amnis ImageStreamX MarkII. The acquired data were analyzed using the IDEAS system.

## ACKNOWLEDGMENTS

This work was supported by grants from the National Natural Science Foundation of China (32270072 to XP, 32200071 to ML, and 82204255 to PZ); the Shandong Provincial Natural Science Foundation (ZR2021QC169 to ML and ZR2023QC172 to YZ); and the Shandong Jianzhu University Domestic Visiting Scholar Program (to ML). The authors thank Xinxing Yang from the University of Science and Technology of China for helpful discussion and the faculty from the Core Facilities for Life and Environmental Sciences at the SKLMT (State Key Laboratory of Microbial Technology, Shandong University) for assistance provided in laser scanning confocal microscopy imaging and flow cytometry analysis.

## References

1. Haeusser DP & Margolin W (2016) Splitsville: structural and functional insights into the dynamic bacterial Z ring. Nature Reviews Microbiology 14(5):305–319.

2. Cameron TA & Margolin W (2024) Insights into the assembly and regulation of the bacterial divisome. Nature Reviews Microbiology 22(1):33–45.

3. Claessen D, Rozen DE, Kuipers OP, Sogaard-Andersen L, & van Wezel GP (2014) Bacterial solutions to multicellularity: a tale of biofilms, filaments and fruiting bodies. Nature Reviews Microbiology 12(2):115–124.

4. Bush MJ, Tschowri N, Schlimpert S, Flardh K, & Buttner MJ (2015) c-di-GMP signalling and the regulation of developmental transitions in streptomycetes. Nat Rev Microbiol 13(12):749–760.

5. Velázquez-Suárez C, et al. (2023) SepT, a novel protein specific to multicellular cyanobacteria, influences peptidoglycan growth and septal nanopore formation in Anabaena sp. PCC 7120. Mbio 14(5).

6. Flardh K & Buttner MJ (2009) *Streptomyces* morphogenetics: dissecting differentiation in a filamentous bacterium. Nat Rev Microbiol 7(1):36–49.

7. Elliot MA, Buttner, M.J., and Nodwell, J.R. (2008) Multicellular development in Streptomyces. *Myxobacteria: Multicellularity and Differentiation* ed Whiteworth DE (American Society for Microbiology), pp 419–438.

8. Schlimpert S, et al. (2017) Two dynamin-like proteins stabilize FtsZ rings during *Streptomyces sporulation*. P Natl Acad Sci USA 114(30):E6176–E6183.

9. Gao S & Hu JJ (2021) Mitochondrial Fusion: The Machineries In and Out. Trends in Cell Biology 31(1):62–74.

10. Bohuszewicz O, Liu JW, & Low HH (2016) Membrane remodelling in bacteria. J Struct Biol 196(1):3–14.

11. Bürmann F, Ebert N, van Baarle S, & Bramkamp M (2011) A bacterial dynamin-like protein mediating nucleotide-independent membrane fusion. Molecular Microbiology 79(5):1294–1304.

12. Junglas B, et al. (2024) Structural basis for GTPase activity and conformational changes of the bacterial dynamin-like protein *Syn*DLP. Cell Rep 43(9).

13. Liu JW, Noel JK, & Low HH (2018) Structural basis for membrane tethering by a bacterial dynamin-like pair. Nature Communications 9.

14. Bush MJ, Gallagher KA, Chandra G, Findlay KC, & Schlimpert S (2022) Hyphal compartmentalization and sporulation in *Streptomyces* require the conserved cell division protein SepX. Nature communication 13(71).

15. Willemse J, Borst JW, de Waal E, Bisseling T, & van Wezel GP (2011) Positive control of cell division: FtsZ is recruited by SsgB during sporulation of *Streptomyces*. Genes Dev 25(1):89–99.

16. Chater K (2011) Differentiation in *Streptomyces*: the properties and programming of diverse cell-types. Streptomyces: Molecular Biology and Biotechnology, ed P D (Caister Academic Press), pp 43–86.

17. Zhang P, et al. (2017) Deletion of MtrA inhibits cellular development of *Streptomyces coelicolor* and alters expression of developmental regulatory genes. Frontiers in microbiology 8:2013.

18. Wang WS, et al. (2013) An Engineered Strong Promoter for *Streptomycetes*. Applied and Environmental Microbiology 79(14):4484–4492.

19. Mccormick JR, Su EP, Driks A, & Losick R (1994) Growth and Viability of *Streptomyces*-*Coelicolor* Mutant for the Cell-Division Gene Ftsz. Molecular Microbiology 14(2):243–254.

20. Sattler L & Graumann PL (2022) Assembly of *Bacillus subtilis* Dynamin into Membrane-Protective Structures in Response to Environmental Stress Is Mediated by Moderate Changes in Dynamics at a Single Molecule Level. Microb Physiol 32(1-2):57–69.

21. Low HH, Sachse C, Amos LA, & Löwe J (2009) Structure of a Bacterial Dynamin-like Protein Lipid Tube Provides a Mechanism For Assembly and Membrane Curving. Cell 139(7):1342–1352.

22. Bianchi-Smiraglia A, et al. (2017) Internally ratiometric fluorescent sensors for evaluation of intracellular GTP levels and distribution. Nat Methods 14(10):1003-+.

23. Presnell KV, Flexer-Harrison M, & Alper HS (2019) Design and synthesis of synthetic UP elements for modulation of gene expression in *Escherichia coli*. Syn Syst Biotechno 4(2):99–106.

24. Rolinson GN (1980) Effect of Beta-Lactam Antibiotics on Bacterial-Cell Growth-Rate. Journal of General Microbiology 120(Oct):317–323.

25. Hopwood DA ed (2007) *Streptomyces in Nature and Medicine* (OXFORD UNIVERSITY PRESS).

26. Deboer PAJ, Crossley RE, & Rothfield LI (1988) Isolation and Properties of Minb, a Complex Genetic-Locus Involved in Correct Placement of the Division Site in Escherichia-Coli. Journal of Bacteriology 170(5):2106–2112.

27. Deboer PAJ, Crossley RE, & Rothfield LI (1992) Roles of Minc and Mind in the Site-Specific Septation Block Mediated by the Mincde System of Escherichia-Coli. Journal of Bacteriology 174(1):63–70.

28. Schwedock J, McCormick JR, Angert ER, Nodwell JR, & Losick R (1997) Assembly of the cell division protein FtsZ into ladder-like structures in the aerial hyphae of Streptomyces coelicolor. Molecular Microbiology 25(5):847–858.

29. Wasserstrom S, et al. (2013) Non-sporulating mutants in *Streptomyces coelicolor* reveal amino acid residues critical for FtsZ polymerization dynamics. Microbiol-Sgm 159:890–901.

30. Wang MF, et al. (2019) Mycobacterial dynamin-like protein IniA mediates membrane fission. Nature Communications 10.

31. Li J, et al. (2018) Impacts of horizontal gene transfer on the compact genome of the clavulanic acid-producing *Streptomyces* strain F613-1. 3 Biotech 8(11).

32. Huang H, Zheng G, Jiang W, Hu H, & Lu Y (2015) One-step high-efficiency CRISPR/Cas9-mediated genome editing in *Streptomyces*. Acta Biochim Biophys Sin (Shanghai) 47(4):231–243.

33. Zhu Y, et al. (2022) The regulatory gene *wblA* is a target of the orphan response regulator OrrA in *Streptomyces coelicolor*. Environmental Microbiology 24(7):3081–3096.

34. Gregory M, Till R, & Smith M (2003) Integration site for *Streptomyces* phage phiBT1 and development of site-specific integrating vectors. J Bacteriol 185(17):5320–5323.

35. Liu M, Xu W, Zhu Y, Cui X, & Pang X (2021) The response regulator MacR and its potential in Improvement of antibiotic production in *Streptomyces coelicolor*. Curr Microbiol 78(10):3696–3707.

36. Karimova G, Pidoux J, Ullmann A, & Ladant D (1998) A bacterial two-hybrid system based on a reconstituted signal transduction pathway. Proceedings of the National Academy of Sciences of the United States of America 95(10):5752–5756.

37. Battesti A & Bouveret E (2012) The bacterial two-hybrid system based on adenylate cyclase reconstitution in Escherichia coli. Methods 58(4):325–334.

