## Supplementary material for "The bifunctional dynamin-like GTPase switch DynAB modulates both vegetative and sporulation cell division in *Streptomyces*": Table S1 and S2

Table S1. Bacterial strains and plasmids used in this study

| Strain or plasmid | Description | Reference |
|---|---|---|
| **Strains** | | |
| *Streptomyces coelicolor* | | |
| M145 | Wild-type, SCP1$^-$ SCP2$^-$ | (1) |
| Δ*mtrA* | *mtrA* mutant strain of M145, Apr$^R$ | (2) |
| Δ*dynA* | unmarked *dynA* deletion strain of M145 | This study |
| Δ*dynB* | unmarked *dynB* deletion strain of M145 | This study |
| Δ*dynAB* | unmarked *dynAB* deletion strain of M145 | This study |
| C-Δ*dynAB* | Δ*dynAB* complemented with *dynAB* of *S. coelicolor* | This study |
| C-Δ*dynAB$_{VEN}$* | Δ*dynAB* complemented with *dynAB* of *S. venezuelae* | This study |
| Δ*ftsZ* | unmarked *ftsZ* deletion strain of M145 | This study |
| Δ*sepX* | unmarked *sepX* deletion strain of M145 | This study |
| Δ*ssgB* | unmarked *ssgB* deletion strain of M145 | This study |
| M145/*dynA** | M145 with *dynA* expressed from the *kasOp** promoter | This study |
| M145/*dynB** | M145 with *dynB* expressed from the *kasOp** promoter | This study |
| M145/*dynAB** | M145 with *dynAB* expressed from the *kasOp** promoter | This study |
| M145/*dynAB$_{VEN}$** | M145 with *dynAB* of *S. venezuelae* expressed from the *kasOp** promoter | This study |
| M145/*ftsZ-egfp** | M145 with the FtsZ-GFP fusion protein gene expressed from the *kasOp** promoter | This study |
| Δ*dynA*/*ftsZ-egfp** | Δ*dynA* with the FtsZ-GFP fusion protein gene expressed from the *kasOp** promoter | This study |
| Δ*dynB*/*ftsZ-egfp** | Δ*dynB* with the FtsZ-GFP fusion protein gene expressed from the *kasOp** promoter | This study |
| Δ*dynAB*/*ftsZ-egfp** | Δ*dynAB* with the FtsZ-GFP fusion protein gene expressed from the *kasOp** promoter | This study |
| M145/*dynAB**/*ftsZ** | M145 with both *dynAB* and *ftsZ* expressed from the *kasOp** promoter | This study |
| M145/*dynAB**/*ftsZ-egfp** | M145 with both *dynAB* and *ftsZ-egfp* expressed from the *kasOp** promoter | This study |
| M145/*sepX-egfp** | M145 with the SepX-GFP fusion protein gene expressed from the *kasOp** promoter | This study |
| Δ*dynAB*/*sepX-egfp** | Δ*dynAB* with the SepX-GFP fusion protein gene expressed from the *kasOp** promoter | This study |
| M145/*dynAB**/*sepX** | M145 with both *dynAB* and *sepX* expressed from the *kasOp** promoter | This study |
| M145/*dynAB**/*sepX-egfp** | M145 with both *dynAB* and *sepX-egfp* expressed from the *kasOp** promoter | This study |
| M145/*egfp-dynAB** | M145 with the GFP-DynAB fusion protein gene | This study |

| | | |
|---|---|---|
| | expressed from the *kasOp** promoter | |
| M145/*dynAB-egfp** | M145 with the DynAB-GFP fusion protein gene expressed from the *kasOp** promoter | This study |
| Δ*ftsZ*/*dynAB-egfp** | Δ*ftsZ* with the DynAB-GFP fusion protein gene expressed from the *kasOp** promoter | This study |
| Δ*sepX*/*dynAB-egfp** | Δ*sepX* with the DynAB-GFP fusion protein gene expressed from the *kasOp** promoter | This study |
| Δ*ssgB*/*dynAB-egfp** | Δ*ssgB* with the DynAB-GFP fusion protein gene expressed from the *kasOp** promoter | This study |
| M145/*ssgB** | M145 with *ssgB* expressed from the *kasOp** promoter | This study |
| Δ*dynAB*/*ssgB** | Δ*dynAB* with *ssgB* expressed from the *kasOp** promoter | This study |
| M145/*dynAB**/*ssgB* * | M145 with both *dynAB* and *ssgB* expressed from the *kasOp** promoter | This study |
| Δ*dynAB*/*dynAB**/*ssgB* * | Δ*dynAB* with both *dynAB* and *ssgB* expressed from the *kasOp** promoter | This study |
| M145/*dyn[AB]* * | M145 with the Dyn[AB] fusion protein gene expressed from the *kasOp** promoter | This study |
| M145/*ssgB**/*dyn[AB]** | M145 with both *ssgB* and *dyn[AB]* expressed from the *kasOp** promoter | This study |
| Δ*dynAB*/*ssgB**/*dyn[AB]** | Δ*dynAB* with both *ssgB* and *dyn[AB]* expressed from the *kasOp** promoter | This study |
| Δ*dynAB*/*dynAB** | Δ*dynAB* with *dynAB*$_{VEN}$ (*S. venezuelae dynAB*) expressed from the *kasOp** promoter | This study |
| Δ*dynAB*/*dynA*$^{K74A}$*B** | Δ*dynAB* with *dynA*$^{K74A}$*B*$_{VEN}$ expressed from the *kasOp** promoter | This study |
| Δ*dynAB*/*dynAB*$^{K129A}$* | Δ*dynAB* with *dynAB*$^{K129A}$$_{VEN}$ expressed from the *kasOp** promoter | This study |
| Δ*dynAB*/*dynA*$^{K74A}$*B*$^{K129A}$* | Δ*dynAB* with *dynA*$^{K74A}$*B*$^{K129A}$$_{VEN}$ expressed from the *kasOp** promoter | This study |
| M145/*dynA*$^{K74A}$*B** | M145 with *dynA*$^{K74A}$*B*$_{VEN}$ expressed from the *kasOp** promoter | This study |
| M145/*dynAB*$^{K129A}$* | M145 with *dynAB*$^{K129A}$$_{VEN}$ expressed from the *kasOp** promoter | This study |
| M145/*dynA*$^{K74A}$*B*$^{K129A}$* | M145 with *dynA*$^{K74A}$*B*$^{K129A}$$_{VEN}$ expressed from the *kasOp** promoter | This study |
| *Escherichia coli* | | |
| DH5α | General cloning strain | Shanghai Weidi |
| DH5α-V | DH5α containing the cloning vector pCE-Zero | This study |
| DH5α/*K-dynAB** | DH5α with *dynAB* expressed from the *kasOp** promoter | This study |
| DH5α/*dynAB-sepX** | DH5α with *dynAB* expressed from the *kasOp** promoter and *sepX* expressed from the *amp*$^R$ | This study |

| | | |
|---|---|---|
| | promoter | |
| DH5α/*dynAB-ssgB** | DH5α with *dynAB* expressed from the *kasOp** promoter and *ssgB* expressed from the *amp*$^{R}$ promoter | This study |
| DH5α/*D-dynAB** | DH5α with *dynAB* expressed from the *rrnD* promoter | This study |
| ET12567(pUZ8002) | Strain used for conjugation between *E. coli* and *Streptomyces* | (1) |
| **Plasmids** | | |
| pMD18-T | General cloning vector | Takara |
| pCE-Zero | General cloning vector | Vazyme |
| pKCcase9d6424 | Shuttle vector for conjugation, Apr$^{R}$ | (3) |
| pMS82 | *Streptomyces* integrative vector, Hyg$^{R}$ | (4) |
| pKC1139 | *Streptomyces/E. coli shuttle* vector, Apr$^{R}$ | (5) |
| pKT25 | Bacterial hybridization vector, Kan$^{R}$ | (6) |
| pUT18c | Bacterial hybridization vector, Amp$^{R}$ | (6) |
| pKNT25 | Bacterial hybridization vector, Kan$^{R}$ | (6) |
| pUT18 | Bacterial hybridization vector, Amp$^{R}$ | (6) |

Abbreviations: Amp$^{R}$, ampicillin resistance; Apr$^{R}$, apramycin resistance; Hyg$^{R}$, hygromycin resistance; Kan$^{R}$, kanamycin resistance

1. Kieser T, Bibb MJ, Buttner MJ, Chater KF, & Hopwood DA eds (2000) *Practical Streptomyces Genetics* (Norwich: John Innes Foundation ), 2nd Edition Ed.
2. Zhang P*, et al.* (2017) Deletion of MtrA inhibits cellular development of *Streptomyces coelicolor* and alters expression of developmental regulatory genes. *Frontiers in microbiology* 8:2013.
3. Huang H, Zheng G, Jiang W, Hu H, & Lu Y (2015) One-step high-efficiency CRISPR/Cas9-mediated genome editing in *Streptomyces*. *Acta Biochim Biophys Sin (Shanghai)* 47(4):231-243.
4. Gregory M, Till R, & Smith M (2003) Integration site for *Streptomyces* phage phiBT1 and development of site-specific integrating vectors. *J Bacteriol* 185(17):5320-5323.
5. Xia HZ & Wang YG (1997) A ketoreductase gene from *Streptomyces mycarofaciens* 1748 DNA involved in biosynthesis of a spore pigment. *Sci China Ser C* 40(6):636-641.
6. Gully D & Bouveret E (2006) A protein network for phospholipid synthesis uncovered by a variant of the tandem affinity purification method in *Escherichia coli*. *Proteomics* 6(1):282-293.

Table S2. Primers sequence (5′→3′) used in this study

| | |
|---|---|
| **For deletion of *dynA*** | |
| *dynA*-sgRNA-F(*Spe*I) | GGATCTTCCAGAGAT**ACTAGT**CTGCTCACCAAGGCAGGCCTGTTTTAGAGCTAGAAATA |
| *dynA*-sgRNA-R | TCAAAAAAAGCACCGACTCGG |
| *dynA*-up-F | CGGTGCTTTTTTTGAGCGCACGATCTACCAGCAC |
| *dynA*-up-R | CGTACGTCCAAGGTCACCAC |
| *dynA*-down-F | GACCTTGGACGTACGGTCGCCACCGTGCTGCTG |
| *dynA*-down-R(*Hind*III) | CTGCCGTTCGACGAT**AAGCTT**CGCCCCAGACTTCCTCCC |
| **For deletion of *dynB*** | |
| *dynB*-sgRNA-F(*Spe*I) | GGATCTTCCAGAGAT**ACTAGT**GACGACCGGGCTGAACAGAAGTTTTAGAGCTAGAAATA |
| *dynB*-sgRNA-R | TCAAAAAAAGCACCGACTCGG |
| *dynB*-up-F | CGGTGCTTTTTTTGACCGCCGACATCTGGGTGATG |
| *dynB*-up-R | CGCCTCCTTGGTCGTGTGC |
| *dynB*-down-F | ACGACCAAGGAGGCGCGCGAGGTGCGGGAACAGT |
| *dynB*-down-R(*Hind*III) | CTGCCGTTCGACGAT**AAGCTT**GCAGAATCAGCAGCCTCACAA |
| **For deletion of *dynAB*** | |
| *dynAB*-sgRNA-F(*Spe*I) | GGATCTTCCAGAGAT**ACTAGT**GACGACCGGGCTGAACAGAAGTTTTAGAGCTAGAAATA |
| *dynAB*-sgRNA-R | TCAAAAAAAGCACCGACTCGG |
| *dynAB*-up-F | CGGTGCTTTTTTTGAGCGCACGATCTACCAGCAC |
| *dynAB*-up-R | CGTACGTCCAAGGTCACCAC |
| *dynAB*-down-F | ACCTTGGACGTACGG CGCGAGGTGCGGGAACAGT |
| *dynAB*-down-R(*Hind*III) | CTGCCGTTCGACGAT**AAGCTT**GCAGAATCAGCAGCCTCACAA |
| **For deletion of *ftsZ*** | |
| *ftsZ*-sgRNA-F(*Spe*I) | GGATCTTCCAGAGAT**ACTAGT**CGATGACTTTGATGACTGCGGTTTTAGAGCTAGAAATA |
| *ftsZ*-sgRNA-R | CTCAAAAAAAGCACCGACTCGG |
| *ftsZ*-up-F | AGTCGGTGCTTTTTTTGAGGACCGACCACCGCCGAG |
| *ftsZ*-up-R | TGATGACTGCGAGGTAGTTCTG |
| *ftsZ*-down-F | CTACCTCGCAGTCATCACGGACTTCCTGAAGTGATAGG |
| *ftsZ*-down-R(*Hind*III) | CTGCCGTTCGACGAT**AAGCTT**GTAACCGACCACGGAACGCA |
| **For deletion of *sepX*** | |
| *sepX*-sgRNA-F(*Spe*I) | GGATCTTCCAGAGAT**ACTAGT**CTGTTCCTTGCCTTCACGCTGTTTTAGAGCTAGAAATA |
| *sepX*-sgRNA-R | CTCAAAAAAAGCACCGACTCGG |
| *sepX*-up-F | GGTGCTTTTTTTGAGGTCAACCTGCCGGGCTCCTC |
| *sepX*-up-R | GTCGAAGAGCGGCGTCCCGACGATGACTGCGTAGATG |
| *sepX*-down-F | ACGCCGCTCTTCGACCAGT |

| | |
|---|---|
| *sepX*-down-R(*Hind*III) | CTGCCGTTCGACGAT**AAGCTT**CCATGCCGTGCAGGAACT |
| **For deletion of *ssgB*** | |
| *ssgB*-sgRNA-F(*Spe*I) | GGATCTTCCAGAGAT**ACTAGT**CGTGTCGTACCGCAGGCCTGGTTTTAGAGCTAGAAATA |
| *ssgB*-sgRNA-R | CTCAAAAAAAGCACCGACTCGG |
| *ssgB*-up-F | AGTCGGTGCTTTTTTTGAGGGGTGCGTCAATGATTCGG |
| *ssgB*-up-R | CACCTACGGTGCCGTTGTA |
| *ssgB*-down-F | ACGGCACCGTAGGTGAGCTAGGGCGGGGCCTCGC |
| *ssgB*-down-R(*Hind*III) | CTGCCGTTCGACGAT**AAGCTT**CCGACCGGAGATTGCGACATT |
| **For verification of unmarked gene deletion** | |
| *dynAB*-confirm-F | CGTGGCTCGATGTGGAGGTTC |
| *dynAB*-confirm-R | CCGTCCAGGCGTTCTTCTCG |
| *ftsZ*-confirm-F | AAGACGTACCCACGCTGGAATT |
| *ftsZ*-confirm-R | CCTGTCGGTGAAGCCGAAGT |
| *sepX*-confirm-F | TGGCGGGACCATCTCGTCTT |
| *sepX*-confirm-R | CGTCAATGTGCGTGGACCTGTG |
| *ssgB*-confirm-F | AAACCTGTCACCGGAATGGG |
| *ssgB*-confirm-R | GTGCCGTATGCGGTTGTCC |
| **For complementation of Δ*dynAB*** | |
| *dynAB*-com-F(*Hind*III) | GAGAACCTAGGATCC**AAGCTT**CGTGGCTCGATGTGGAGGTTC |
| *dynAB*-com-R(*Kpn*I) | TGAAAAACGCTCACT**GGTACC**CCGTCCAGGCGTTCTTCTCG |
| *dynAB*$_{sven}$-com-F(*Hind*III) | GAGAACCTAGGATCC**AAGCTT**GGACTCGAACCTGCGGCCAAGTGC |
| *dynAB*$_{sven}$ -com-R(*Kpn*I) | TGAAAAACGCTCACT**GGTACC**AAACGGAACCTCGCCACCC |
| **For expression of *Streptomyces* genes** | |
| *kasOp* *-F(*Kpn*I) | GGATCTTCCAGAGAT**GGTACC**GTTCACATTCGAACCGTC |
| *kasOp* *-R | AACTCCCCCAGTCCTGCACG |
| HX-*dynA*-F | AGGACTGGGGGAGTTGTGGTGACCTTGGACGTACGG |
| HX-*dynA*-R(*Hind*III) | CTGCCGTTCGACGAT**AAGCTT**TCACCTCTCCTTCTGCAGTAC |
| HX-*dynB*-F | AGGACTGGGGGAGTTGTGACCGCCGTCACTGACCAG |
| HX-*dynB*-R(*Hind*III) | CTGCCGTTCGACGAT**AAGCTT**CTACCTCGTCGCCGTCCCCGCCGTTG |
| HX-*dynA*$_{sven}$-F | AGGACTGGGGGAGTTGTGGTGACCTTGGACGAACG |
| HX-*dynB*$_{sven}$-R(*Hind*III) | CTGCCGTTCGACGAT**AAGCTT**CTACCTGCCCCGTACCCCCGTC |
| *kasop**-F(*Spe*I) | TCGTTAGTTAGGCTA**ACTAGT**TGTTCACATTCGAACCGTC |
| HX-*ftsZ*-F | AGGACTGGGGGAGTTATGGCAGCACCGCAGAACTACCT |

| | |
|---|---|
| HX-*ftsZ*-R(*Eco*RV) | CATGATTACGAATTC**GATATC**CTACTTCAGGAAGTCCGG CACGTCCA |
| HX-*sepX*-F | AGGACTGGGGGAGTTATGAGCAGCAGCGGCCTCATCTA |
| HX-*sepX*-R(*Eco*RV) | CATGATTACGAATTC**GATATC**CTACTCGTTGGCCGCGCG GGGGCGGTC |
| HX-*ssgB*-F | AGGACTGGGGGAGTTGTGGCATGTCGATTTCGCCGAC |
| HX-*ssgB*-R(*Eco*RV) | CATGATTACGAATTC**GATATC**CTAGCTTTCCGCCAGGAT GTG |
| **For expression of GFP-tagged proteins** | |
| *kasOp* *-F(*Kpn*I) | GGATCTTCCAGAGAT**GGTACC**GTTCACATTCGAACCGTC |
| *kasOp* *-R | AACTCCCCCAGTCCTGCACG |
| N-*egfp*-F | AGGACTGGGGGAGTTATGGTGAGCAAGGGCGAGGA |
| N-*egfp*-R | CCGCCAGAGCCACCTCCGCCTGAACCGCCTCCACCCTT GTACAGCTCGTCCATGC |
| *egfp*-*dynAB*-F | GCGGAGGTGGCTCTGGCGGTGGCGGTAGTGTGACCTTG GACGTACGG |
| *egfp*-*dynAB*-R(*Hind*III) | CTGCCGTTCGACGAT**AAGCTT**ATCGATCTACCTCGTCGC CGTCCCCGCCGTTG |
| *dynAB*-*egfp*-F | AGGACTGGGGGAGTTGTGGTGACCTTGGACGTACGGGT GACCTTGGACGTACGG |
| *dynAB*-*egfp*-R | CCAGAGCCACCTCCGCCTGAACCGCCTCCACCCCTCGT CGCCGTCCCCGCCGTTG |
| C-*egfp*-F | TCAGGCGGAGGTGGCTCTGGCGGTGGCGGTAGTATGGT GAGCAAGGGCGAGGA |
| C-*egfp*-R(*Hind*III) | CTGCCGTTCGACGAT**AAGCTT**TCACTTGTACAGCTCGTC CATGC |
| *kasop**-F(*Spe*I) | TCGTTAGTTAGGCTA**ACTAGT**TGTTCACATTCGAACCGT C |
| *ftsZ*-*egfp*-F | AGGACTGGGGGAGTTATGGCAGCACCGCAGAACTACCT |
| *ftsZ*-*egfp*-R | CTGAACCGCCTCCACCCTTCAGGAAGTCCGGCACGTCC A |
| *sepX*-*egfp*-F | AGGACTGGGGGAGTTATGAGCAGCAGCGGCCTCATCTA |
| *sepX*-*egfp*-R | CTGAACCGCCTCCACCCTCGTTGGCCGCGCGGGGGC |
| C-*egfp*-R(*Eco*RV) | CATGATTACGAATTC**GATATC**TCACTTGTACAGCTCGTCC AT |
| **For expression of fusion protein Dyn[AB]** | |
| *kasOp* * F (pMS) | TGAAAAACGCTCACTTGTTCACATTCGAACCGTCTCTGC |
| DynA(without TGA)-R | AGTGACGGCGGTCACCCTCTCCTTCTGCAGTACGGA |
| DynB-F | GTGACCGCCGTCACTGACCAGGAC |
| DynB-R(pMS) | AGCCGAGAACCTAGGATCCCTACCTCGTCGCCGTCCCCG CCGTTGT |
| **For mutation of GTP binding P-loop domain in DynAB$_{ven}$** | |
| *kasOp* * F (pMS) | TGAAAAACGCTCACTTGTTCACATTCGAACCGTCTCTGC |
| DynA$_{sven}$ (K74A) R | TTGACGAGCGTGGAAGCGCCGGCTCCGGTGGA |

| | |
|---|---|
| DynA*sven*(K74A) F | GCTTCCACGCTCGTCAACTCCCTTGTCGGG |
| DynB*sven*(K129A) R | GGTGGAGGCGCCGCTGCCGGTGGCGCCGGCGATCGCGATGACGGTGT |
| DynB*sven*(K129A) F | AGCGGCGCCTCCACCCTCTTCAACGCCCTGGCCGGCGTCCCCGTCTCCGAG |
| DynB*sven* R(pMS) | AGCCGAGAACCTAGGATCCCTACCTGCCCCGTACCCCCGT |
| **For bacterial hybridization experiments** | |
| *ssgB pkt25-F* | CGGGCTGCAGGGTCGACTCTAGAAATGGCATGTCGATTTCGCCGAC |
| *ssgB pkt25-R* | TGCTCGAGGTCGACGGTATCGATTCAGCTTTCCGCCAGGATGTGCG |
| *ssgB Put18c-F* | CGCCACTGCAGGTCGACTCTAGAGATGGCATGTCGATTTCGCCGAC |
| *ssgB Put18c-R* | CCATATTACTTAGTTATATCGATTCATCAGCTTTCCGCCAGGATGTGCG |
| *sepX put18 F* | CAGGAAACAGCTATGAGCAGCAGCGGCCTCATC |
| *speXput18 R* | GGGATCCTCTAGAGTCTCGTTGGCCGCGCGGGGGCGGTC |
| *sepH put18 F* | CAGGAAACAGCTATGCCCGAACTGCGTGTCGTGG |
| *sepH put18 R* | GGGATCCTCTAGAGTCTCCTGCTTCTTGCGCCGCGTG |
| *dynA put18 F* | CAGGAAACAGCTATGGTGACCTTGGACGTACGGC |
| *dynA put18 R* | GGGATCCTCTAGAGTCCTCTCCTTCTGCAGTACGG |
| *dynB put18 F* | CAGGAAACAGCTATGACCGCCGTCACTGACCAGGAC |
| *dynB put18 R* | GGGATCCTCTAGAGTCCTCGTCGCCGTCCCCGCCGTT |
| *pkc1139-prrnD-F* | TATGACATGATTACGAATTC**GATATC**CAGAAAAAAAGATCAAAAAAA |
| SsgB-pkc1139-R | AGCTTGGGCTGCAGGTCGACTCTAGACTAGCTTTCCGCCAGGATGT |
| *pkc1139-prrnD-F* | TATGACATGATTACGAATTCGATATCCAGAAAAAAAGATCAAAAAAA |
| SepX-pkc1139-R | AGCTTGGGCTGCAGGTCGACTCTAGACTACTCGTTGGCCGCGCGGGGGC |
| *pkc1139-prrnD-F* | TATGACATGATTACGAATTCGATATCCAGAAAAAAAGATCAAAAAAA |
| *DynA-pkc1139-R* | AGCTTGGGCTGCAGGTCGACTCTAGATCACCTCTCCTTCTGCAGTACGGA |
| **For expression of genes in *E. coli*** | |
| *prrnD*-F(*Eco*RV) | GGATCTTCCAGAGATCAGAAAAAAAGATCAAAAAAATAC |

| | |
|---|---|
| *prrnD*-R | CACCATAGCTGTTTCCTGTGTGAAATTGTTATTCGTCTCA ACGGAGGCGCATTA |
| *prrnD-DynAB*$_{sven}$-F | AACAGCTATGGTGACCTTGGACGAACG |
| *prrnD-DynAB*$_{sven}$-R | CTGCCGTTCGACGATCTACCTGCCCCGTACCCCCGTC |
| *kasOp**-F(*Kpn*I) | GGATCTTCCAGAGAT**GGTACC**GTTCACATTCGAACCGTC |
| *kasOp**-R | AACTCCCCCAGTCCTGCACG |
| HX-*dynA*$_{sven}$-F | AGGACTGGGGGAGTTGTGGTGACCTTGGACGAACG |
| HX-*dynB*$_{sven}$-R(*Hind*III) | CTGCCGTTCGACGAT**AAGCTT**CTACCTGCCCCGTACCCC CGTC |
| *pAmp*$^{R}$-F (*Spe*I) | ACCCTCACTAAAGGG**ACTAGT**CTGACGCTCAGTGGAAC GAA |
| *pAmp*$^{R}$-R | CAGTCGATCATAGCACGATC |
| *pAmp*$^{R}$-*sepX*$_{sven}$-F | TGCTATGATCGACTGATGAGCAGCAGCGGCCTCATC |
| *pAmp*$^{R}$-*sepX*$_{sven}$-R | GTTTAAACCTGCAGGTCATTCGTTGGCCGCCCGGG |
| *pAmp*$^{R}$-*ssgB*$_{sven}$-F | TGCTATGATCGACTGATGAACACCACGGTCAGCTG |
| *pAmp*$^{R}$-*ssgB*$_{sven}$-R | GTTTAAACCTGCAGGTCAGCTGTCGGCCAGGATGTG |
| **For Real-time PCR analysis** | |
| *sco1541*-RT-F | GCATGTCGATTTCGCCGAC |
| *sco1541*-RT-R | CAGCTGACCGTGGTGTTCAT |
| *sco2685*-RT-F | CCGACATCTGGGTGATGGTC |
| *sco2685*-RT-R | GGTGGCGTCGTACTCCTTC |
| *sco2684*-RT-F | GGACTGGTCCTGATCGACCT |
| *sco2684*-RT-R | GATAGCGCTCGTGGAGTACG |
