## Supplementary material for "The bifunctional dynamin-like GTPase switch DynAB modulates both vegetative and sporulation cell division in *Streptomyces*": Figure Supplementary

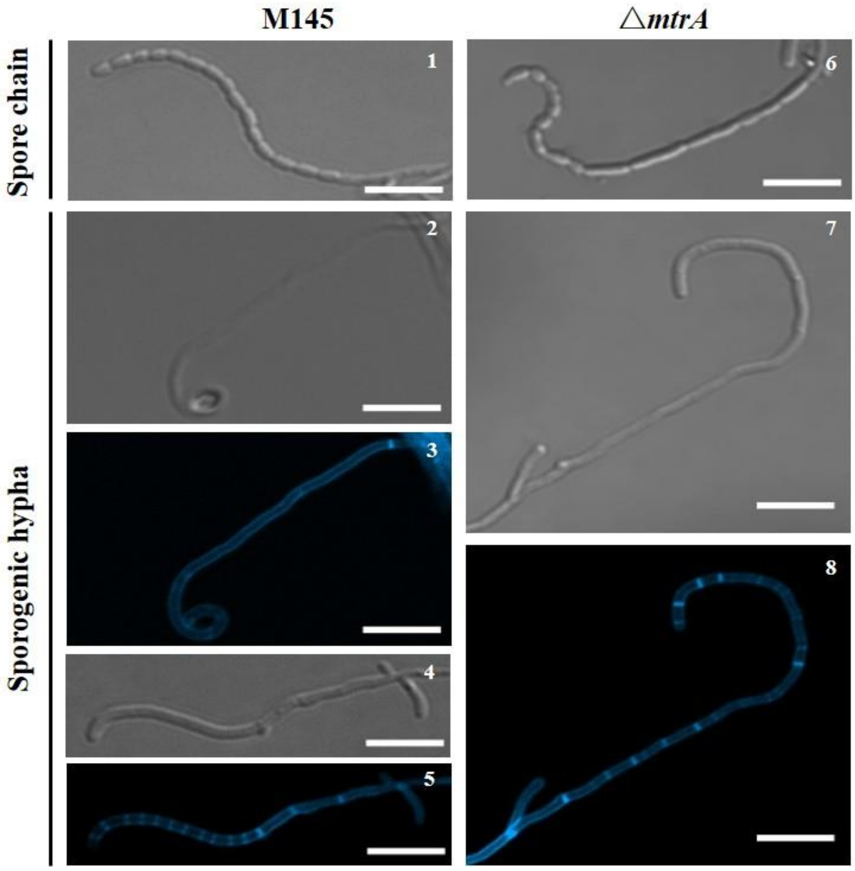


Fig. S1. Impaired cell division in an *mtrA* mutant. Light and fluorescence micrographs of the *S. coelicolor* wild-type strain M145 and Δ*mtrA* grown on BSCA agar supplemented with HADA, showing the irregular spores and sporulation septa formed in the sporogenic hyphae of Δ*mtrA*. Scale bar, 5 μm.

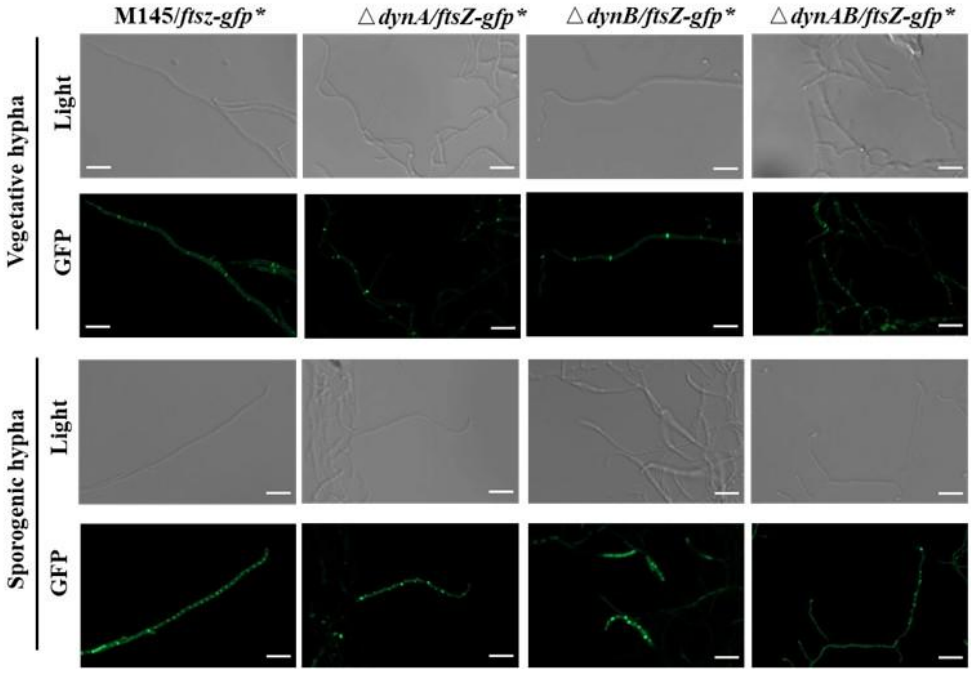


Fig. S2. Impact of DynA, DynB, and DynAB on localization of FtsZ in *S. coelicolor*. Gene *ftsZ*, fused with *gfp* at the 3′ end, was expressed from the *kasO* promoter. *Streptomyces* strains expressing GFP-tagged FtsZ were grown on HADA-supplemented BSCA agar. The light images show the vegetative hyphae and sporogenic hyphae of the resulting strains, and the fluorescence images show the impaired FtsZ-rings formed in the vegetative hyphae and sporogenic hyphae of Δ*dynA* and Δ*dynB*, compared with the typical Z-ring formed in M145. Scale bar, 5 μm.

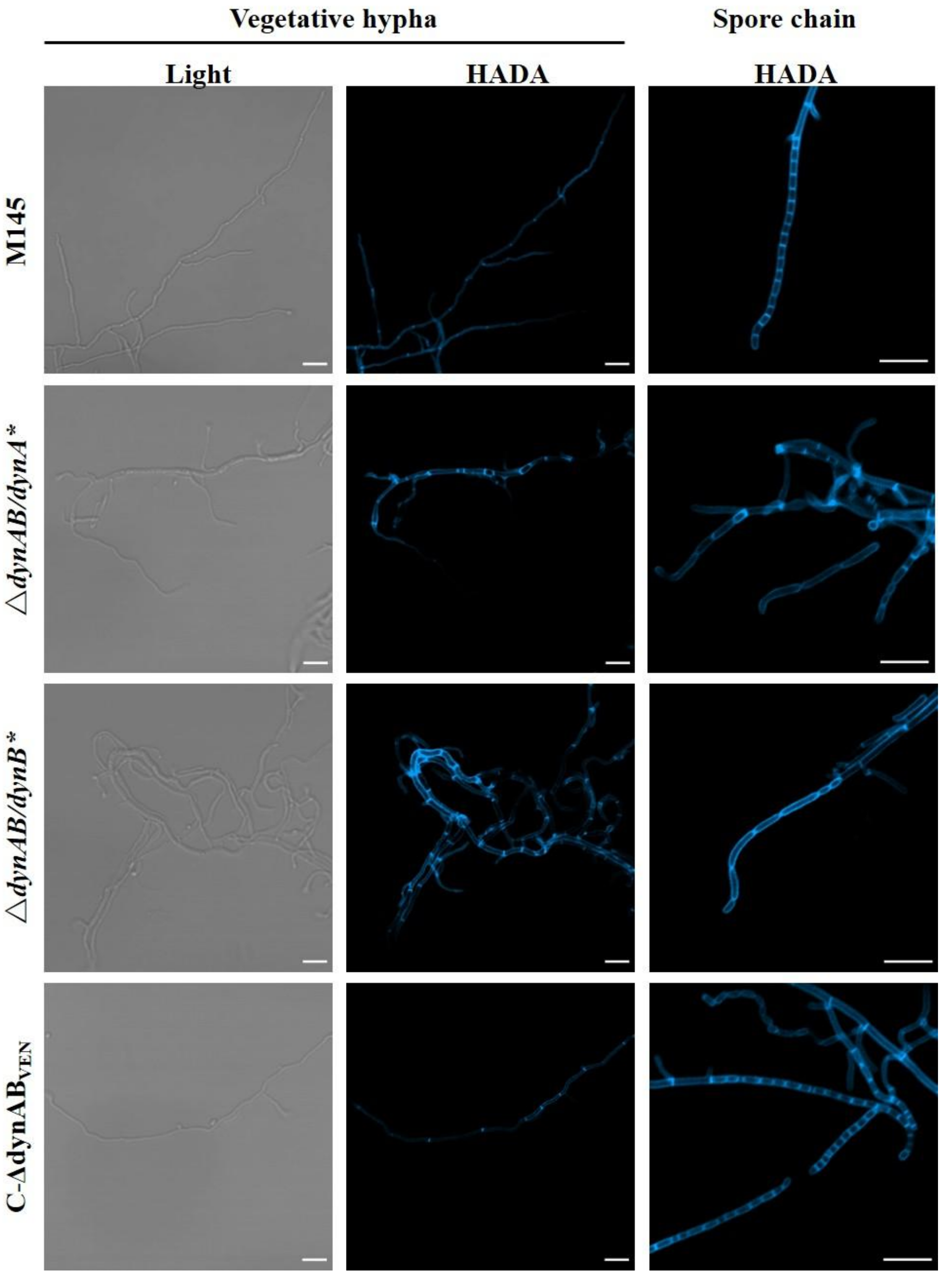


Fig. S3. Complementation of *S. coelicolor* Δ*dynAB* with *S. coelicolor dynA* or *dynB* or *S. venezuelae dynAB*. *S. coelicolor* genes *dynA* and *dynB* were expressed from the *kasO* promoter, and *S. venezuelae dynAB* were expressed from their native promoter. M145 and its derivative strains were grown on HADA-supplemented BSCA agar. The light images show the vegetative hyphae, and the fluorescence images show the cross-walls formed in the vegetative hyphae and sporulation septa formed in the sporogenic hyphae. Scale bar, 5 µm.

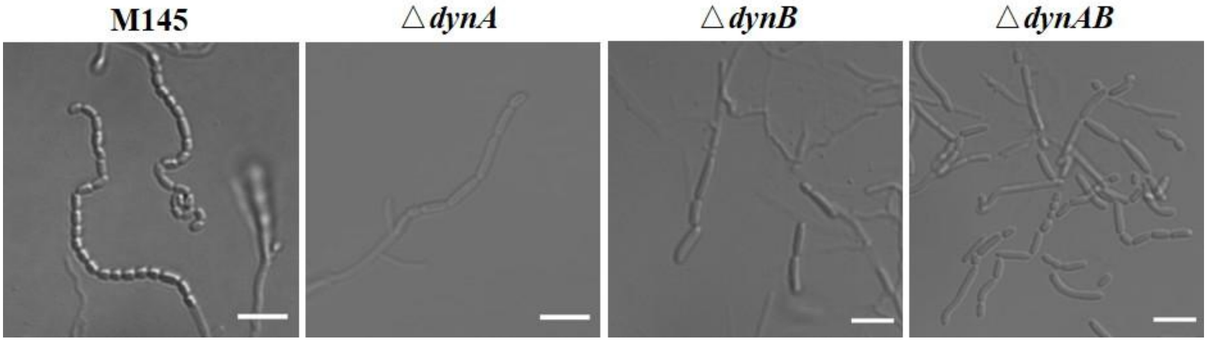


Fig. S4. Impact of DynAB on sporulation-specific cell division. M145, Δ*dynA*, Δ*dynB*, and Δ*dynAB* were grown on BSCA agar. Images show the irregular spore chains formed in Δ*dynA*, Δ*dynB*, and Δ*dynAB*, compared with in M145, and indicate that deletion of DynA, DynB, or DynA and DynB impaired sporulation-specific cell division in *S. coelicolor*. Scale bar, 5 μm.

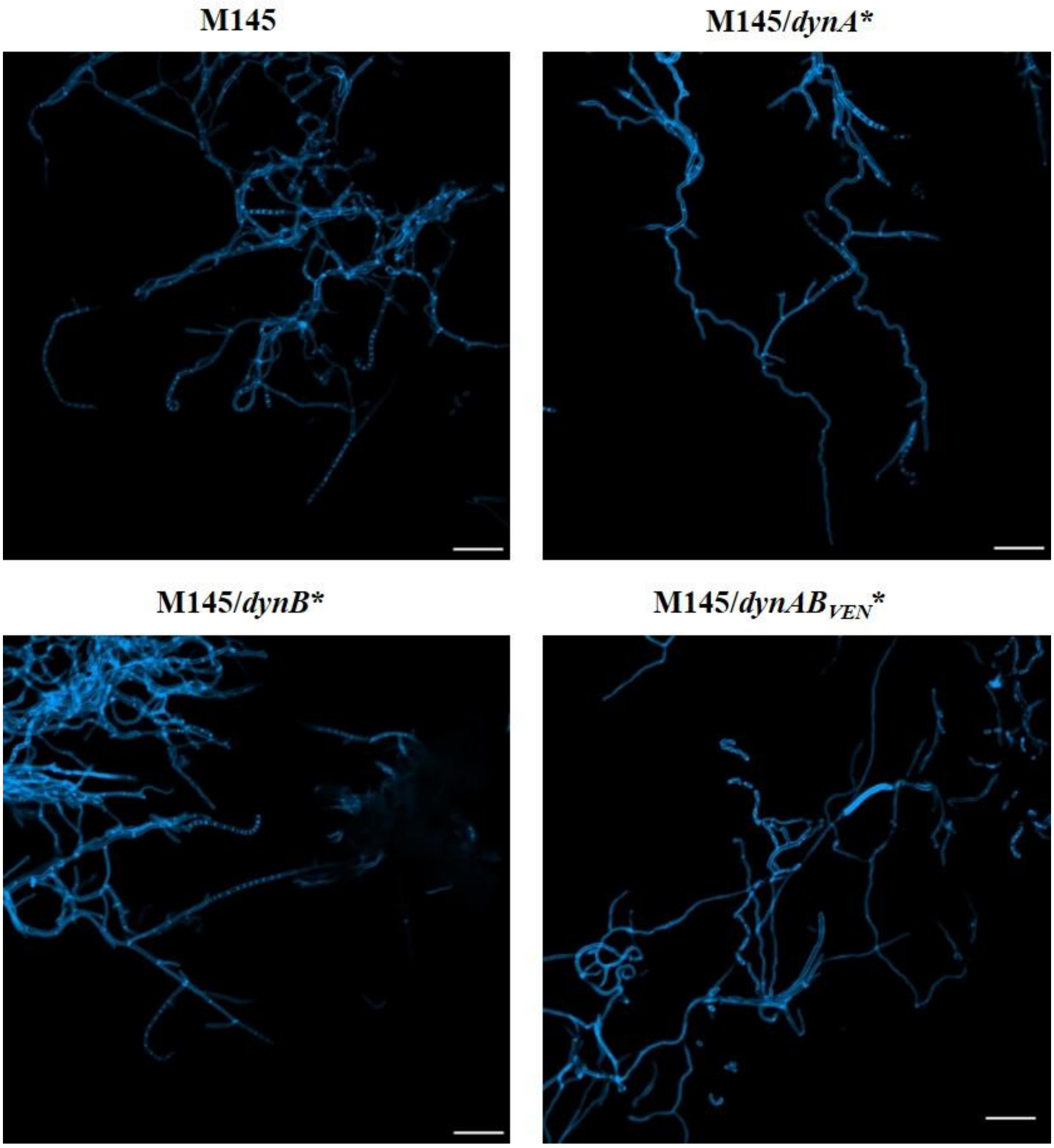


Fig. S5. Expression of DynA or DynB alone does not repress cross-wall formation in *S. coelicolor*. M145 strains constitutively expressing *dynA*, *dynB*, or *dynAB* (dynABsco*) of *S. coelicolor* and *dynAB* ($dynAB_{VEN}$*) of *S. venezuelae*. were grown on HADA-supplemented BSCA agar. The images show the cross-walls formed during vegetative growth, with only the vegetative hyphae of M145/$dynAB_{SVE}$* lacking cross-walls. Scale bar, 5 µm.

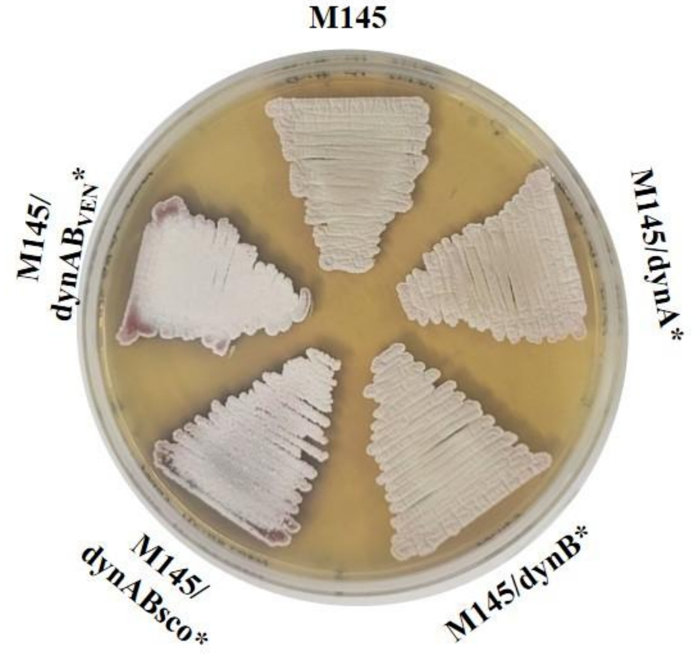


Fig. S6. Phenotypes of M145 strains constitutively expressing *dynA*, *dynB*, or *dynAB* (dynABsco*) of *S. coelicolor* and *dynAB* (dynAB$_{VEN}$*) of *S. venezuelae*. The photograph was taken after 72 hours of growth on BSCA.

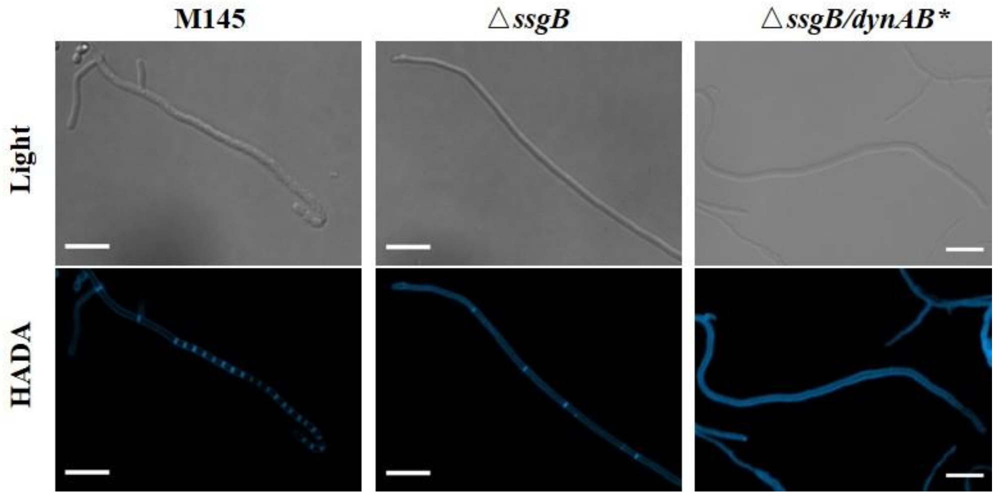


Fig. S7. DynAB repress cross-wall-like structures in the sporogenic hyphae of Δ*ssgB*. Deletion of *ssgB* leads to formation of cross-wall-like septa in the sporogenic hyphae of *S. coelicolor*. Cross-wall-like septa were present in Δ*ssgB* whereas this phenotype was completely repressed by expressing DynAB in Δ*ssgB*. M145, Δ*ssgB*, and Δ*ssgB*/*dynAB** were grown on HADA-supplemented BSCA agar. The light images showed the sporogenic hyphae, and the fluorescence images show the cross-walls in the vegetative hyphae and cross-wall-like septa in the sporogenic hyphae. Scale bar, 5 μm.

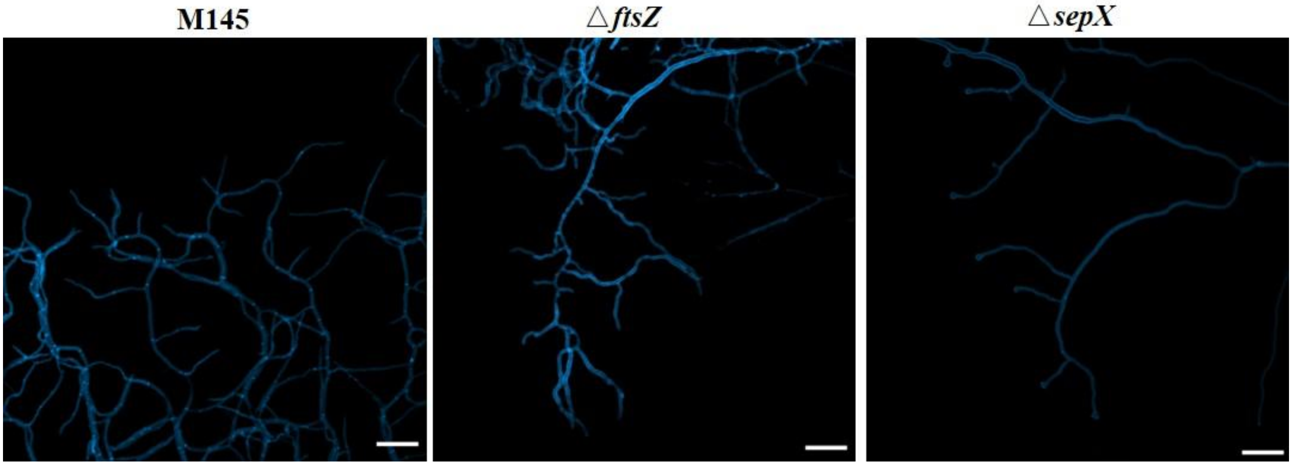


Fig. S8. FtsZ and SepX are essential for cross-wall formation in *S. coelicolor*. M145, Δ*ftsZ*, and Δ*sepX* were grown on HADA-supplemented BSCA agar. The images show that no cross-walls were formed in the vegetative hyphae of Δ*ftsZ* and Δ*sepX* whereas cross-walls were detected in the parental strain M145, during vegetative growth. Scale bar, 5 μm.

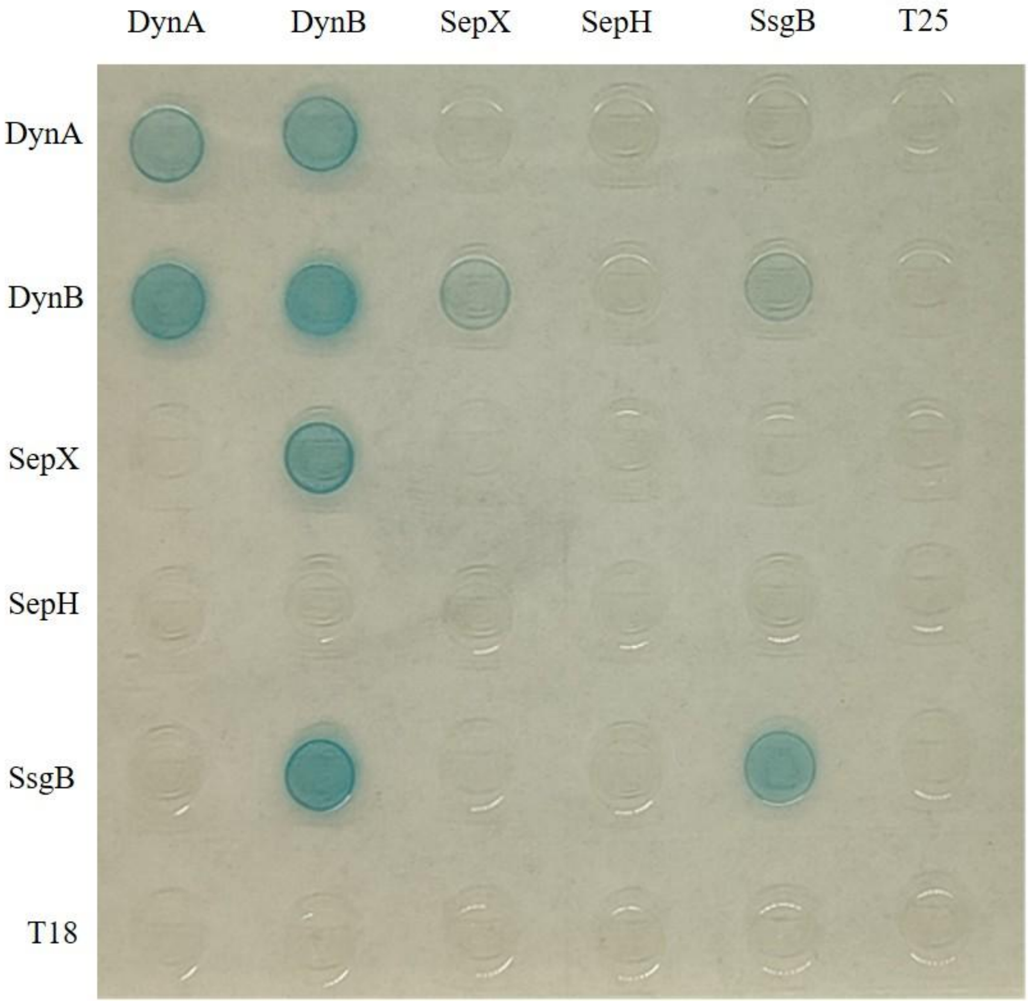


Fig. S9. Bacterial two-hybrid interaction assays of DynA and DynB with other cell division proteins of *S. coelicolor*. Direct, pairwise interactions were examined among DynAB and SepX, SepH, and SsgB constructs based on pKT25 and pUT18C. The pKT25 and pUT18C vectors were used as negative controls. Each combination of compatible plasmids was co-transformed into *E. coli* strain BTH101, and the resulting strains with fusion or control plasmids were induced with the addition of IPTG and inoculated on LB plates with IPTG and X-gal. The blue color results from restoration of adenylate cyclase activity and indicates protein-protein interactions between the two fusion proteins.

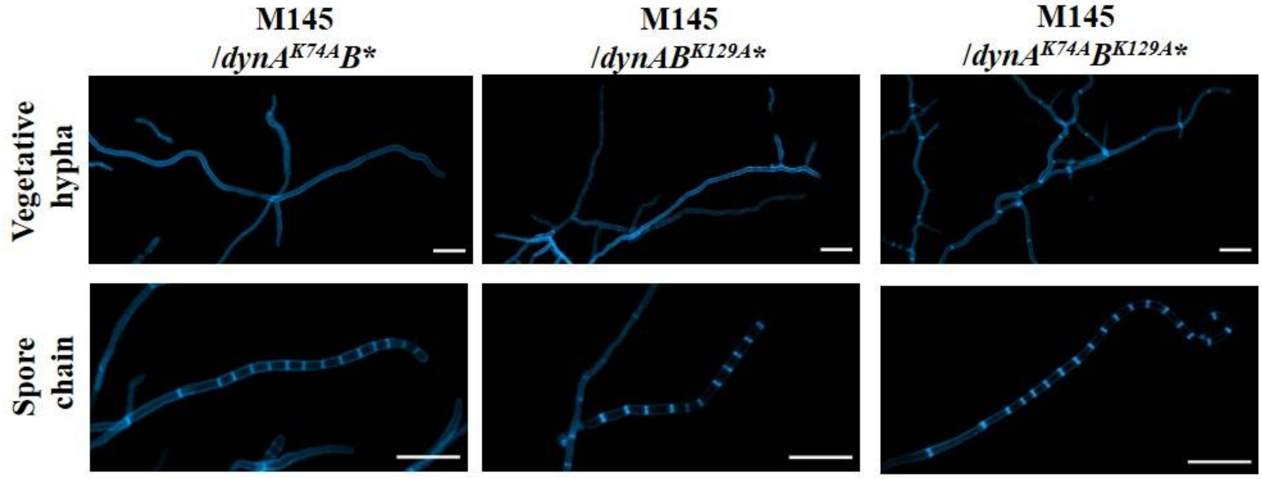


Fig. S10. Functional analysis of DynAB with mutated GTP-binding domains in M145. The phenotype of M145 strains expressing DynA(K74A), DynB(K129A), or both DynA(K74A) and DynB(K129A) was examined following growth on HADA-supplemented BSCA agar. The blue fluorescence images show the position of cross-walls in the vegetative hyphae or sporulation septa in the aerial hyphae. Cross-walls were not formed in strains with mutation of a single GTP-binding domain, but were formed in the strain with mutations in both domains (M145/d*ynA*$^{K74A}$*B*$^{K129A}$*). Scale bar, 5 µm.

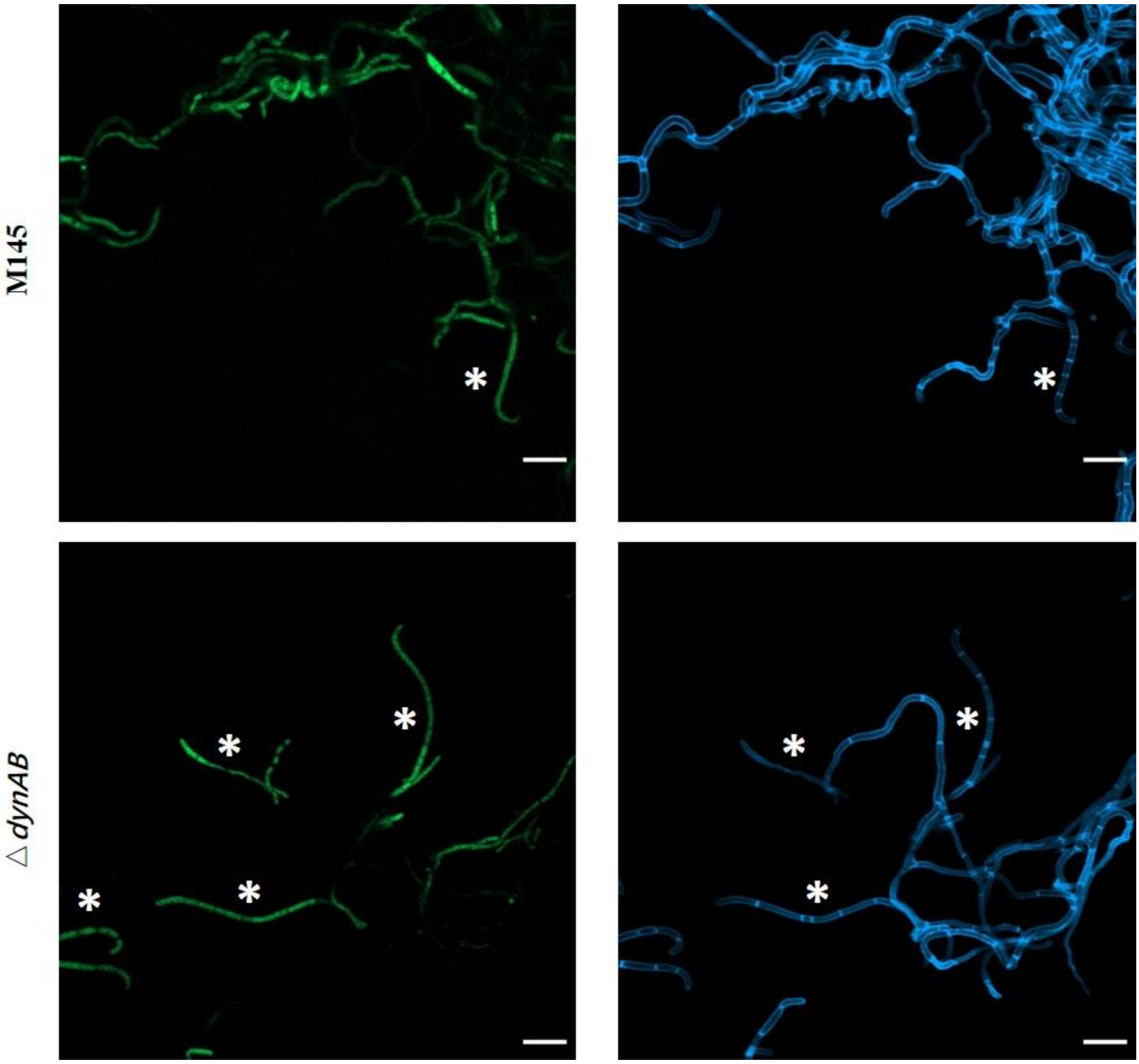


Fig. S11. GTP levels in different types of *S. coelicolor* cells. The green fluorescence images show the cellular level of GFP, determined using the GTP-sensing protein GEVAL30. The blue fluorescence images show the position of cross-walls in the vegetative hyphae or sporulation septa in the sporogenic hyphae. These images demonstrated that the level of GFP was generally higher in sporogenic hyphae. Asterisks indicate sporogenic hyphae. Scale bar, 5 µm.

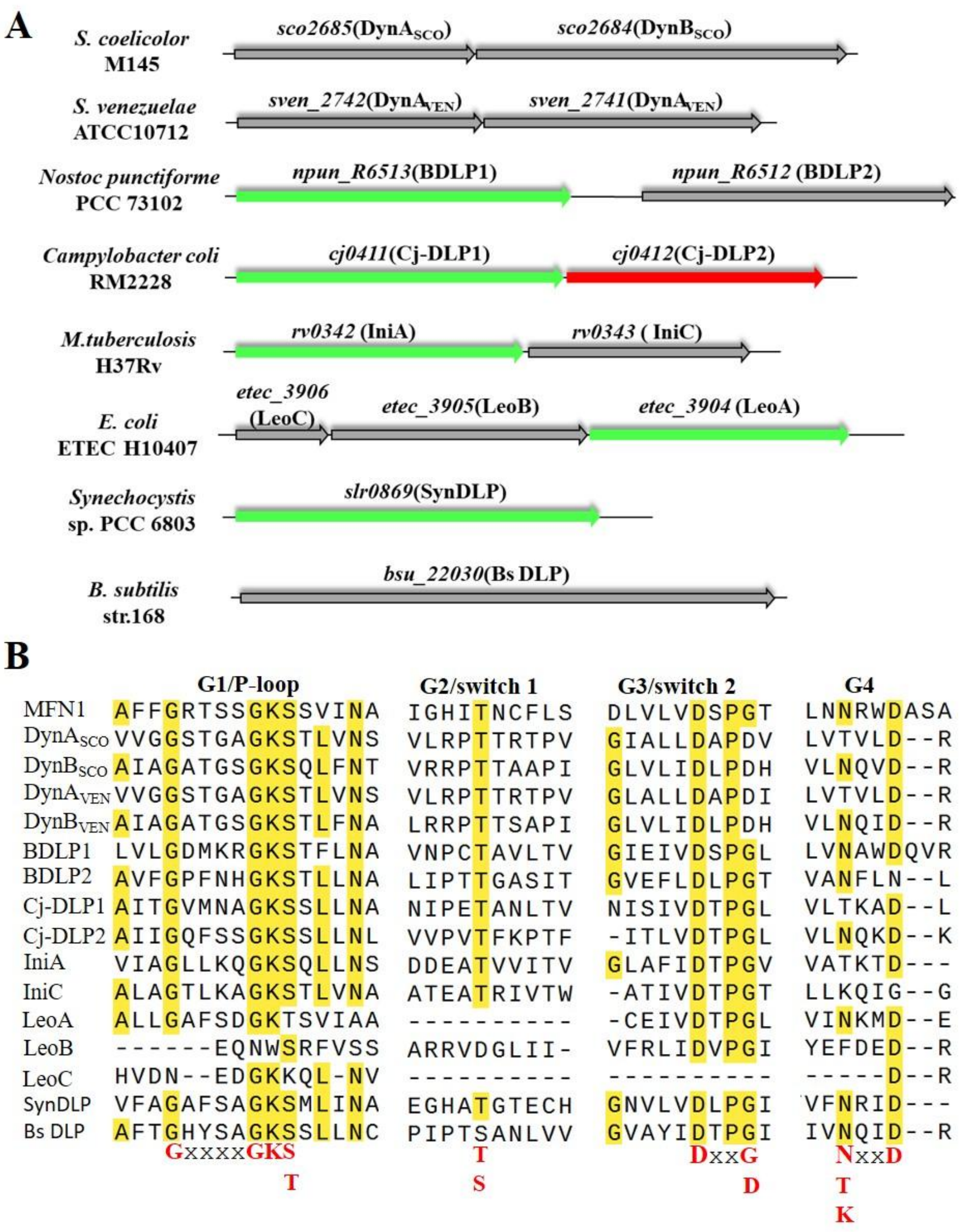


Fig. S12. Similarity of DynAB and other conserved bacterial DLP pairs. (A) Genetic organization of *dynAB* and homologous genes in other bacteria. The BDLP genes for Cj-DLP1 and Cj-DLP2 in *Campylobacter coli*, BDLP1 and BDLP2 in *Nostoc punctiforme*, and *iniA* and *iniC* in *Mycobacterium tuberculosis* are arranged tandemly. In *E. coli*, the first gene became divided into two genes (LeoC and LeoB), forming a three gene operon, whereas only one BDLP gene is present, potentially due to the fusion of two genes, in a *Synechocystis* species (SynDLP) and *Bacillus subtilis* (DynA). (B) Alignment of the G1-G4 GTP binding motifs of DynA and DynB within the dynamin family. Human MNF1 (UniProt accession Q8IWA4) serves as a representative for eukaryotic DLPs in this alignment. The yellow or red color depicts proteins with resolved structure, and the grey color depicts proteins whose structure has not been resolved yet.

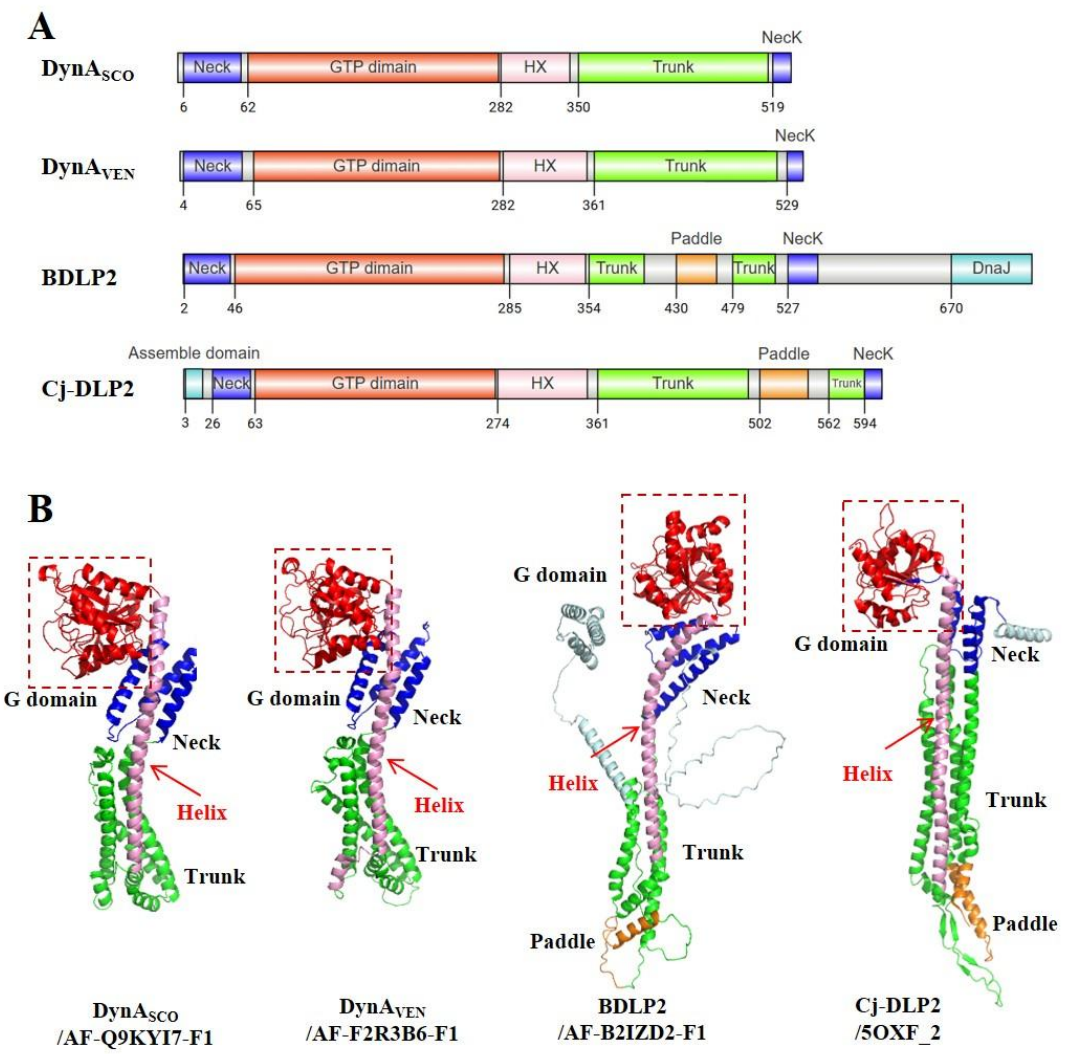


Fig. S13. Structural analysis of DynA. (A) Domain organization in DynA and homologous proteins. Domains with structural homology and/or functional equivalence are depicted in the same color. The helices linking the GTP, neck, and trunk domains are highlighted in pink. The numbers indicate the amino acid position in the primary structures of these DLPs. (B) Predicted structure of DynA of *S. coelicolor* and *S. venezuelae*, BLDP2, and the resolved structure of Cj-DLP2. The structure identification codes following the protein names were obtained from the Protein Data Bank (PDB) or from the AlphaFold Protein Structure Database.

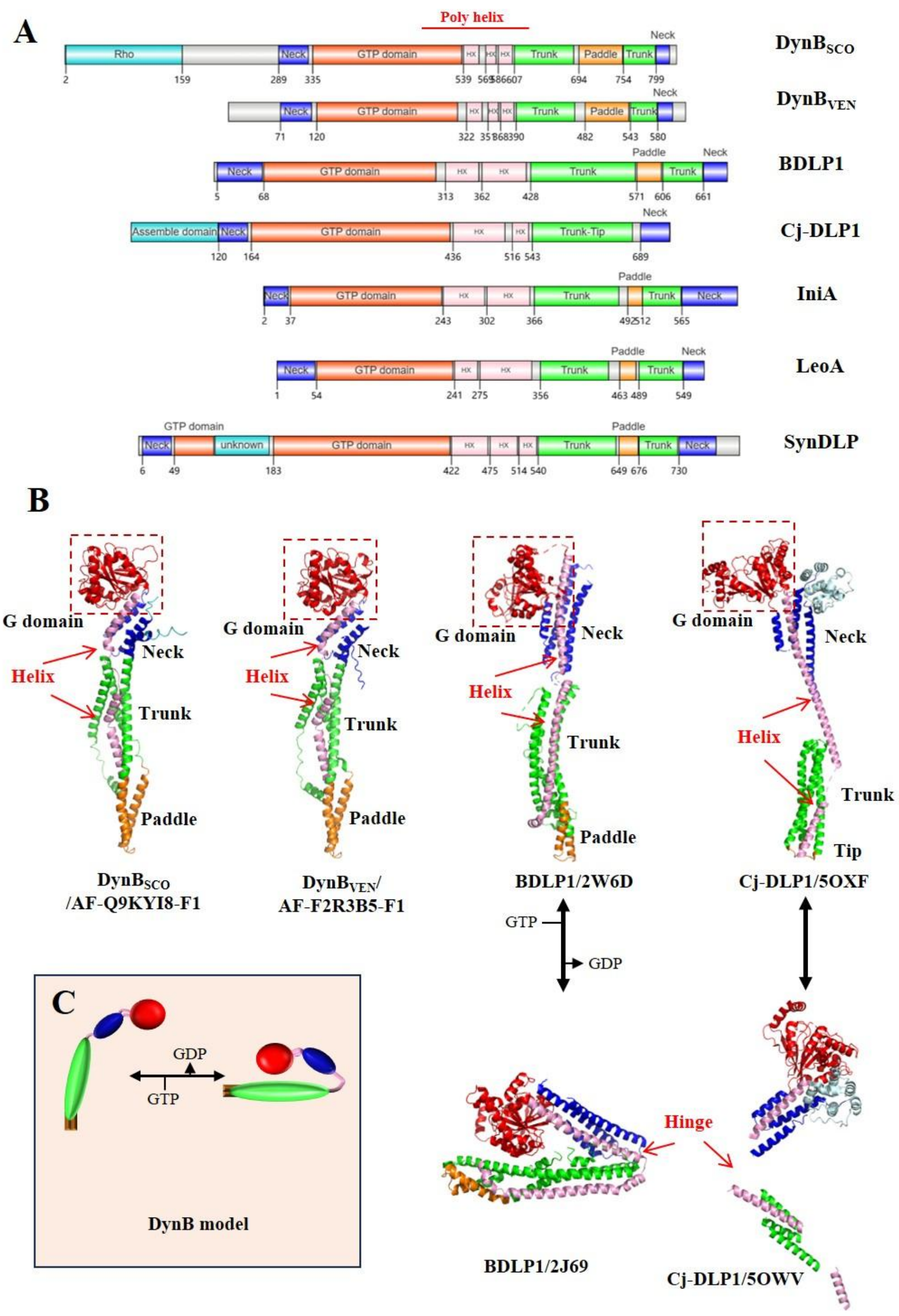


Fig. S14. Structural analysis of DynB. (A) Domain organization in DynB and homologous proteins. Domains with structural homology and/or functional equivalence are depicted in the same color. The helices linking the GTP, neck, and trunk domains are highlighted in pink. The numbers indicate the amino acid position in the primary structures of these DLPs. (B) Predicted structure of DynB of *S. coelicolor* and *S. venezuelae*, and the resolved structure of BLDP1 and Cj-DLP1. The structure identification codes following the protein names were obtained from the Protein Data Bank (PDB) or from the AlphaFold Protein Structure Database. (C) BDLP1 and Cj-DLP1 may undergo conformational changes upon hydrolysis of GTP. The similarity between the predicted structures of DynB, BDLP1, and Cj-DLP1 suggests that the poly-helix structure connecting the GTP, neck, and trunk domains may allow for significant conformational changes.
